# Structural basis of polyspecific drug recognition by MRP2

**DOI:** 10.64898/2026.05.26.728010

**Authors:** Naito Ishimoto, Hailey Kadonoff, Eleanor J. McGuigan, Lina Yousof, Theodoros I Roumeliotis, Jyoti S Choudhary, Kenneth J. Linton, Konstantinos Beis

**Affiliations:** Department of Life Sciences, Imperial College London, SW7 2AZ, London, United Kingdom; Blizard Institute, Faculty of Medicine and Dentistry, Queen Mary University of London, E1 2A, London, UK; Functional Proteomics group, Chester Beatty Laboratories, The Institute of Cancer Research, London, SW3 6JB, UK

## Abstract

Multidrug resistance-associated protein 2 (MRP2/ABCC2) is a hepatobiliary ATP-binding cassette transporter that mediates the efflux of endogenous conjugates and many therapeutic agents, yet the basis of its broad ligand specificity remains poorly understood. Here we combine cryo-electron microscopy, ATPase measurements and cell-based transport inhibition assays to define how MRP2 recognizes clinically relevant compounds from distinct chemical classes. We find that chemically diverse ligands occupy overlapping regions of a shared transmembrane cavity but adopt different poses, contact networks and stoichiometries. Ligand recognition is mediated predominantly by a permissive environment complemented by ligand-specific polar interactions, rather than by a single conserved pharmacophore. These findings establish a structural framework for MRP2 polyspecificity and provide insight into hepatobiliary drug disposition and transporter-mediated drug-drug interactions.

## Introduction

Multidrug resistance-associated protein 2 (MRP2/ABCC2) is the only C-type ATP-binding cassette (ABC) transporter expressed in the apical (canalicular) membrane of hepatocytes where it is a major mediator of the biliary excretion of endogenous metabolites, drug conjugates and therapeutic compounds^1^. MRP2 is expressed to a lesser extent in enterocytes, renal proximal tubule cells and placental syncytiotrophoblasts, and thus contributes to excretory and barrier functions at multiple tissue interfaces. In doing so, it shapes drug disposition, drug-drug interactions and, as an off-target of drug inhibition, susceptibility to transporter-mediated toxicity.

MRP2 is a polyspecific anion transporter that actively effluxes metabolites such as bilirubin-glucuronide (inhibition or mutation of MRP2 presents primarily as hyperbilirubinaemia), estradiol-17β-D-glucuronide, sulphated bile acids, leukotrienes and a wide range of clinical drugs, including bosentan, vinblastine, cisplatin, methotrexate and their Phase II conjugates ^2,3^. Accordingly, the clearance of many clinical agents depends, at least in part, on MRP2. Substrate classes include chemotherapeutics of many distinct chemical classes; pteridines, methotrexate, anthracyclines (doxorubicin, epirubicin), camptothecin derivatives (SN-38-glucuronide), vinca alkaloids, platinum drugs, monomethylsulfonates (HIV protease inhibitors such as saquinavir, lopinavir, ritonavir), nucleotide analogues (adefovir, cidofovir), carboxylic acid esters (statins such as pravastatin), monocarboxylic acids (angiotensin receptor blockers such as valsartan, olmesartan), several β-lactam and cephalosporin antibiotics, macrolides, and dietary or herbal constituents such as flavonoids and silybin. This broad specificity underpins its dual role in systemic clearance and in local protection of the liver, intestinal epithelia and placenta from xenobiotic accumulation^1^.

The physiological importance of MRP2 is illustrated by Dubin-Johnson syndrome, a hereditary conjugated-hyperbilirubinaemia caused by biallelic loss-of-function variants that impair canalicular export of bilirubin glucuronides and related organic anions^4,5^. More common regulatory and coding variants have also been associated with altered exposure or toxicity for several drugs, including methotrexate, mycophenolic acid, statins, calcineurin inhibitors and antiretrovirals. Together with its susceptibility to inhibition by co-administered compounds, these observations establish MRP2 as an important determinant of inter-individual variability in drug handling.

We have previously determined the structure of *Rattus norvegicus* Mrp2 (rMrp2), a mammalian homologue with 78% identity plus 11% similarity to human MRP2, by cryo-EM in apo and drug (probenecid) bound states^6^. The overall architecture of rMrp2 is composed of a small N-terminal transmembrane domain (TMD0) that associates with two further TMDs, TMD1 and TMD2. TMD0 is directly linked to TMD1 by the lasso linker (L_0_). TMD1 and TMD2 each consist of 6 TM helices and form the substrate/drug binding site. They are linked to two nucleotide binding domains (NBDs), NBD1 and NBD2, where ATP binding and hydrolysis powers the transport cycle. The apo structure of rMrp2 revealed an autoinhibited conformation in which the unphosphorylated regulatory (R)-domain which links NBD1 to TMD2 was localised to the interface between the two transmembrane domains blocking access to both the substrate/drug binding pocket and transition of the protein to an outward open conformation. A similar R-domain conformation was observed in the human MRP2 cryo-EM structure^7^. Transport assays with rMrp2 reconstituted in proteoliposomes showed that the fully phosphorylated protein had increased transport activity compared to the partially phosphorylated or fully dephosphorylated protein^6^. Regulation by phosphorylation was further confirmed in a physiologically-relevant *in vitro* model of human tissue^6^. We also determined the cryo-EM structure of rMrp2 with probenecid (a drug used for the treatment of gout but jaundice is a side effect)^8,6^. Our probenecid bound structure provided insights into the modulation of rMrp2 transport activity by drug-drug interactions. Probenecid is an MRP2 transport substrate^3^ and it can also stimulate the transport of other compounds such as the chemotherapy drugs paclitaxel or methotrexate^9,3^. The drug-bound structure revealed two probenecid molecules bound within the TMD binding pocket. The TMDs of the MRP class of transporters is typically characterised by a positively charged P-pocket and a hydrophobic H-pocket^10^. The probenecid bound structure identified two additional drug binding sites; binding site 1 shares some overlap with the P-pocket whereas the second site is distal to both H-/P-pockets and drug binding site ^16^. We have also shown that transport of the fluorescent substrate 5(6)-Carboxy-2′,7′-DichloroFluorescein (CDF) is inhibited by probenecid in proteoliposomes^6^, so probenecid can act as both a stimulator and competitive inhibitor of the transport of other compounds. The probenecid-bound structure indicated that rMrp2 can support multi-ligand binding, suggesting a structural basis for substrate-dependent inhibition or stimulation.

Despite its physiological and pharmacological importance, the molecular basis of MRP2 polyspecificity remains poorly understood. MRP2-interacting compounds differ markedly in scaffold, size, flexibility, charge distribution and hydrophobicity, making transporter recognition difficult to predict from chemical class alone (Figure 1 and Supplementary Figure 1). Here we explore the nature of this unusual polyspecificity by determining the cryo-EM structure of rMrp2 in complex with several clinically relevant drugs and inhibitors from distinct chemical classes. Surprisingly, these chemically diverse ligands have overlapping binding pockets despite lacking obvious shared chemical or physicochemical determinants, revealing an unexpectedly adaptable TMD that can also accommodate multiple ligand stoichiometries.

**Figure 1.**
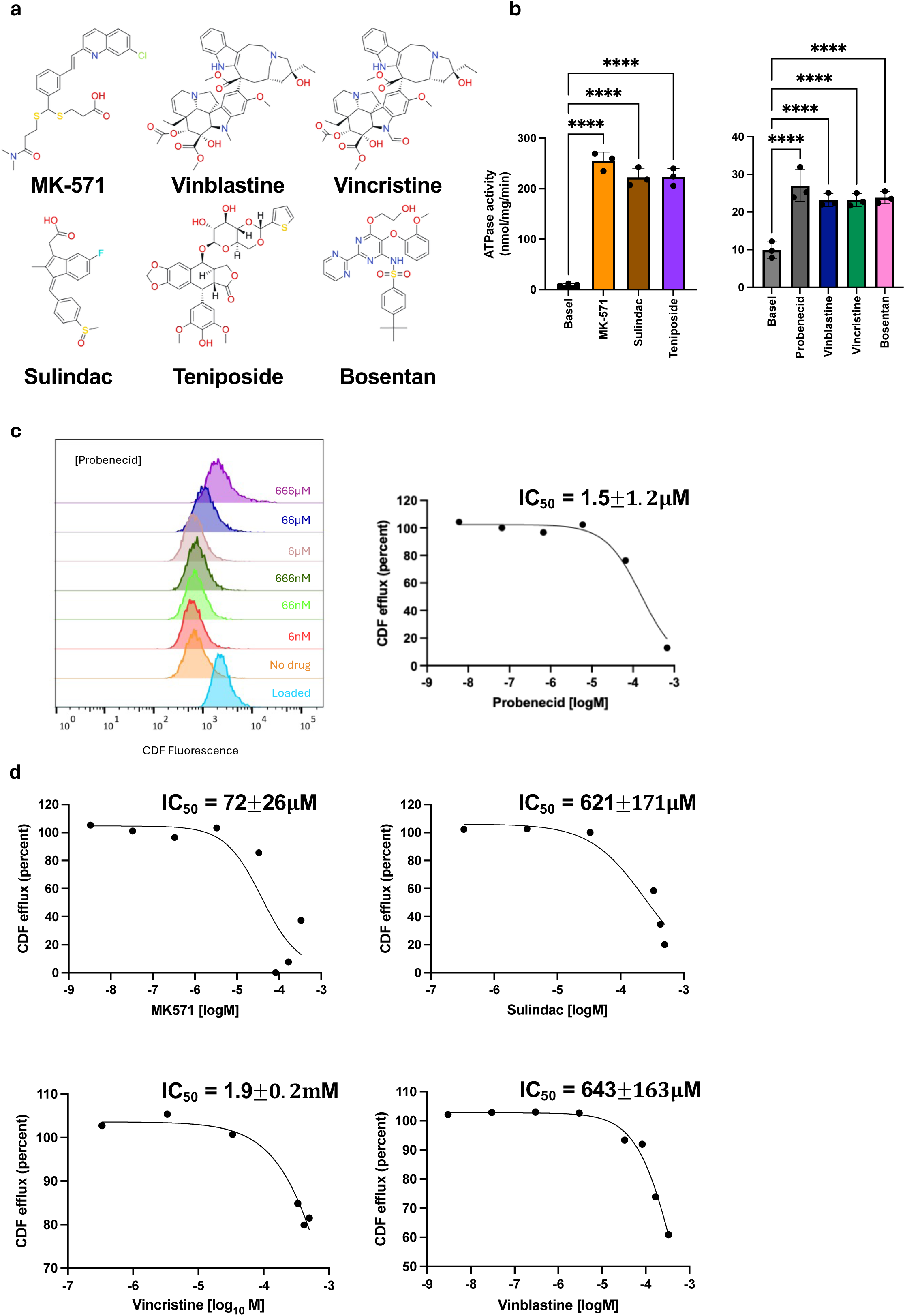
Polyspecificity of rMrp2. (a) Chemical structures of the drugs and inhibitors that are recognised by MRP2. (b) All tested drugs stimulate rMrp2 ATPase activity to varying degrees. MK-571, sulindac, and teniposide display nearly 30-fold greater stimulation compared to the other drugs, which exhibit lower activity. (c) Determination of IC□□ values for inhibition of CDF efflux from loaded cells mediated by rMrp2. Flow cytometric histograms showing inhibition of CDF efflux by probenecid in HEK293-FlpIn-rMrp2 cells. The ‘loaded’ sample represents cells following incubation with CDFDA. The ‘no drug’ sample represents maximal CDF efflux from the loaded cells. The remaining six samples were incubated with increasing concentrations of probenecid as indicated. Median CDF fluorescence for each population was subtracted from the median of the ‘loaded’ sample to quantify CDF efflux. Percent efflux at each concentration was then plotted to determine the half maximal inhibitory concentration. IC_50_ represented as the mean +/- SEM (n=3). (d) IC_50_ values for four additional compounds were calculated using the same method as described for probenecid.

ABC transporters are common off-targets of chemotherapeutic agents and the ensuing drug-induced liver injury remains an important challenge in drug development and clinical practice^11,12^. MRP2 has been proposed to contribute to resistance against some chemotherapeutic agents, we therefore examined whether transporter expression is sufficient to alter drug response in cells. Stable expression of rMrp2 in the human epithelial kidney cell line HEK293, failed to confer resistance to the cytotoxic drugs vinblastine and vincristine. Consistent with this, our analysis of public proteomic and drug-response datasets across human cancer cell lines did not reveal a clear association between MRP2 abundance and sensitivity to common anticancer drugs. However, 5 of the 6 drugs for which we present structural data did inhibit the efflux of CDF by rMrp2. These structural and functional data *in cellulo* demonstrate that each of four distinct classes of drugs can competitively inhibit rMrp2 function.

Our findings provide a structural framework for understanding MRP2 polyspecificity and competitive inhibition, showing that chemically diverse compounds can engage overlapping sites within a broadly permissive binding pocket. The cellular data further suggest that ligand binding and inhibition of transport do not necessarily translate into measurable resistance to cytotoxic drugs. Accordingly, in the systems examined here, our results do not support a major role for MRP2 as a direct mediator of resistance to vinca alkaloids, vinblastine and vincristine, but remain consistent with a role in drug-transporter interactions that may influence drug disposition and drug-induced liver injury.

## Results

### Functional characterisation of distinct chemotypes that interact with rMrp2

We have established a robust protocol for the expression and purification of rMrp2, a tractable homologue for biochemical and structural studies^6^. Within the substrate-binding pocket rMrp2 shares 98% sequence identity with the human transporter, enabling direct functional and mechanistic comparisons (Supplementary Figure 2). To investigate the drug polyspecificity of rMrp2, we assembled a panel of clinically used compounds with diverse pharmacological activities and distinct chemotypes that are established MRP2 substrates (Figure 1a and Supplementary Table 1). This panel included teniposide (an epipodophyllotoxin glycoside), vincristine and vinblastine (vinca alkaloids), sulindac (an arylalkanoic acid non□steroidal anti□inflammatory drug (NSAID)), probenecid (a sulfonamide uricosuric agent), bosentan (a diaryl□sulfonamide) and the experimental compound MK□571 (a heteroaryl□sulfonamide). We evaluated their interaction with rMrp2 by measuring drug□stimulated ATP hydrolysis. All compounds stimulated the basal ATPase activity of rMrp2; teniposide, sulindac and MK-571 stimulated by nearly 30-fold (Figure 1b). These data are consistent with broad ligand recognition by rMrp2 and supported subsequent structural and functional analysis.

To functionally assess drug interaction with rMrp2, we performed rMrp2 inhibition assays in HEK293-FlpIn cells stably expressing rMrp2 or carrying an empty vector (see Materials and Methods) (Supplementary Figure 3). Transport activity was quantified by monitoring the cellular efflux of CDF, a well-characterised substrate of rMrp2^6^. We previously demonstrated that purified rMrp2, when reconstituted into proteoliposomes, can transport CDF in an ATP-dependent manner, confirming that CDF transport is a direct readout of rMrp2 activity^6^. Briefly, cells were loaded with 20 *μ*M 5(6)-carboxy-2’,7’-dichlorofluorescein diacetate (CDFDA). Once inside the cells, CDFDA is metabolically de-esterified to the fluorescent compound CDF, which is membrane-impermeant and therefore requires active transport to be exported (Figure 1c). After loading, cells were exposed to increasing concentrations of the different test drugs, and the decrease in intracellular CDF fluorescence was used as a measure of rMrp2-mediated efflux. In control experiments, HEK293-FlpIn-Mrp2 cells expressing rMrp2 showed robust CDF efflux, whereas HEK293-FlpIn cells carrying an empty vector exhibited minimal CDF export, confirming that CDF transport in this system is mediated by the recombinant rMrp2. Under these conditions, five of the six drugs tested, reduced CDF efflux in a concentration-dependent manner, consistent with competition for rMrp2-mediated transport (Figure 1c and d).

MK-571 exhibited the highest apparent affinity for rMrp2, with an IC_50_ value of 72 ± 26 *μ*M. Probenecid showed markedly lower inhibition, with a low millimolar IC_50_ value of 1.5 ± 1.2 mM, indicating that it is also a competitor of CDF transport, and in agreement with our previous studies. Vincristine and vinblastine are structurally similar (a methyl group in vinblastine is replaced by a formyl group in vincristine), displayed a weaker inhibitory effect compared to MK-571, with IC_50_ of 1.9 ± 0.2 mM and 0.64 ± 0.16 mM, respectively. This statistically significant difference (unpaired t-test; P = 0.0082, n = 3) suggests that subtle chemical modifications may influence recognition and transport by rMrp2 (. Although there is no distinction between the two drugs in stimulating the ATPase activity of rMrp2 (both drugs induce a 2.5-fold ATPase stimulation). Sulindac also acted as a relatively weak inhibitor, with an IC_50_ of 0.62 ± 0.17 mM, consistent with a lower affinity for the transporter. Pairwise statistical comparison of all measured IC50 valuse are presented in Supplementary Table 2. We were unable to determine an IC_50_ value for teniposide (a potent stimulator of the ATPase activity) under our experimental conditions, as it precipitated at higher concentrations and induced rapid cell death, which precluded reliable quantification of CDF efflux.

### Structural analysis of distinct clinical drugs in complex with rMrp2

#### *Vinca alkaloids* - vincristine and vinblastine

Based on the functional data, we performed structural studies for all the tested drugs to understand the ability of rMrp2 to recognise the different drug classes. We examined how rMrp2 recognises the closely related vinca alkaloids vincristine and vinblastine. Vincristine and vinblastine are complex asymmetric indole alkaloids characterised by a vindoline and catharanthine moiety. The main structural difference between the two compounds is in the tertiary amine of the vindoline moiety that carries a methyl group in vinblastine and a formyl group in vincristine.

We determined the cryo-EM structure of vincristine and vinblastine at 3.46 Å and 3.47 Å resolution (Figure 2, 3 and Supplementary Figure 4 and 5), respectively. Density was well resolved for the core transporter in both complexes, whereas density for TMD0 was weaker, consistent with disorder in this domain. Inspection of the rMrp2-vincristine TMD revealed strong coulomb density for two vincristine molecules (Figure 2a), one within the H-pocket and the other at the interface of drug binding sites 1 and 2 (Figure 2b). The two vincristine molecules display intermolecular interactions with amino acid side chains in the rMrp2 cavity and with each other. The vincristine molecule located between drug binding pocket and 1 and 2 makes hydrogen bonds with Met594 and Arg1253 and extensive van der Waals interactions with the TMD (Figure 2c). The molecule in the H-pocket also makes extensive van der Waals interactions with the TMD and hydrogen bonds to Arg1096 and Arg1253 (Figure 2c). The drug-to-drug interaction is mediated by the formyl and the carboxylic acid methyl ester groups in the vindoline moiety from one drug and the carboxylic acid methyl ester and hydroxyl group of the catharanthine moiety of the other. The presence of two vincristine molecules has been reported in the cryo-EM structures of MRP1^13^, an Mrp2 homologue. In the MRP1 structure, the first vincristine molecule, which is associated with a GSH molecule, is in the drug binding pocket 1 and 2, but displays a different binding pose to the rMrp2 structure, and the second vincristine (without GSH) is found in the translocation pathway (Supplementary Figure 6); the authors concluded that the second molecule is likely not physiologically relevant as it does not display any direct interactions with the transporter^13^.

**Figure 2.**
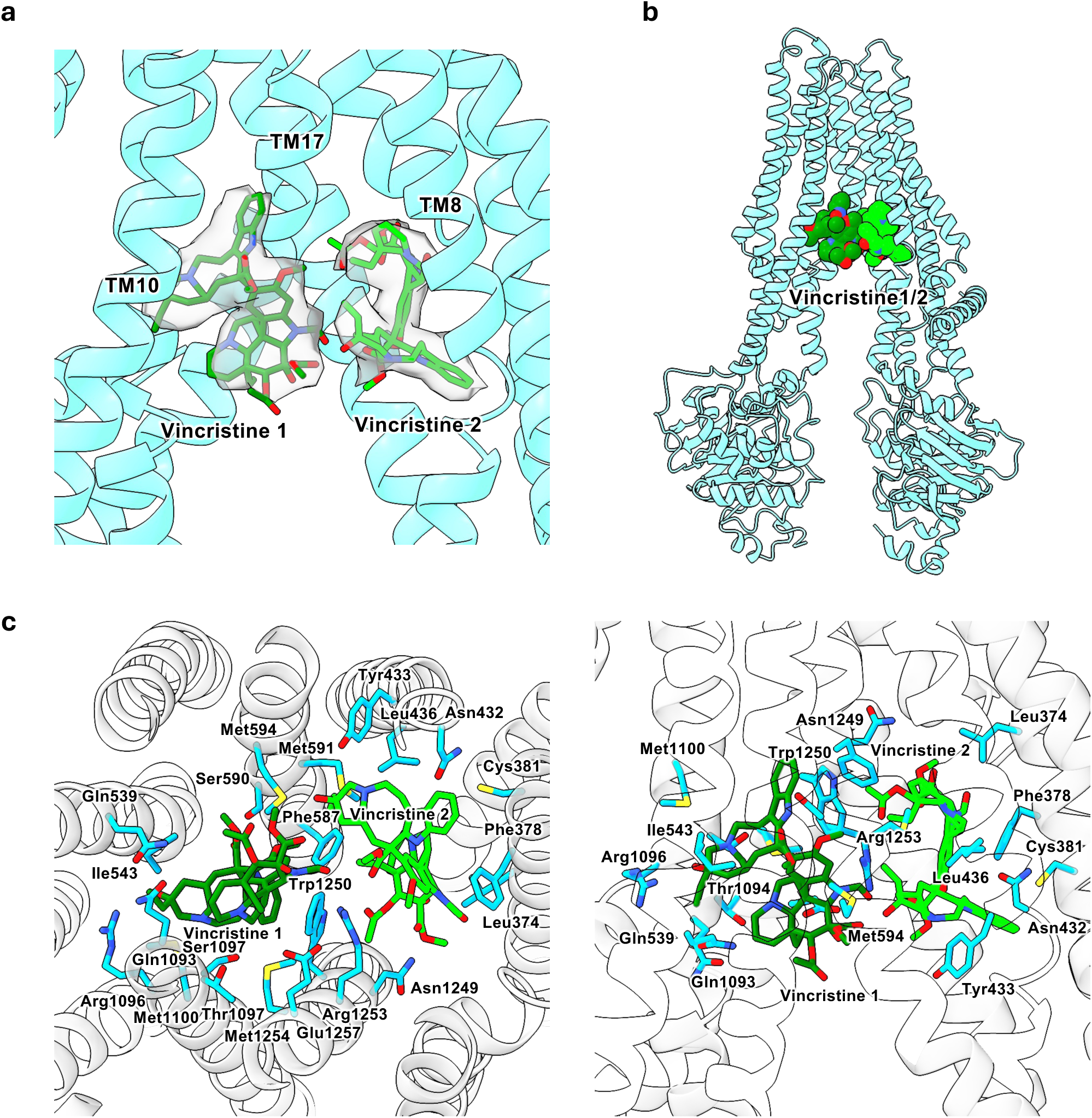
Structural basis of vincristine recognition by rMrp2. (a) Coulomb density (grey surface) and fitted model of two vincristine molecules bound within the TMD. The two vincristine molecules are shown in dark and light green, respectively. (b) Overall structure of the rMrp2 in complex with vincristine. (c) Close-up view of the drug binding site from two orientations. Coordinating residues are shown as cyan sticks.

**Figure 3.**
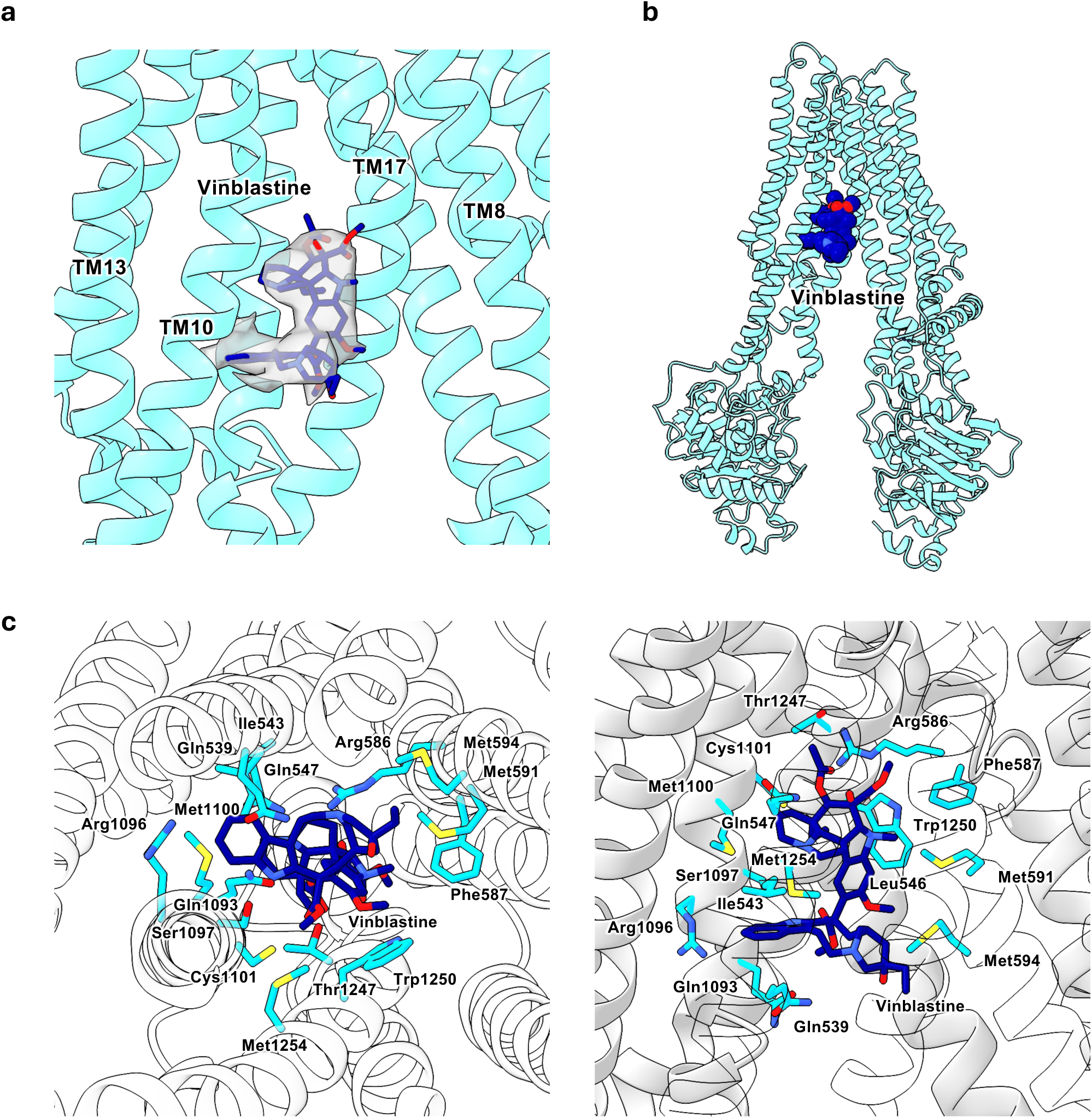
Structural basis of vinblastine recognition by rMrp2. (a) Coulomb density for vinblastine within the TMD, shown as a grey surface, with the fitted ligand model in dark blue. (b) Overall structure of rMrp2 in the vinblastine-bound state. (c) Close-up views of the binding pocket from two orientations, with coordinating residues shown as cyan sticks.

Inspection of the rMrp2-vinblastine maps revealed the presence of density for one molecule of vinblastine, unlike the two bound vincristine molecules in rMrp2-vincristine (Figure 3a). The vinblastine molecule overlaps partly with the H-pocket and translocation pathway (Figure 3b). The vindoline moiety is stabilised by hydrogen bonds with Arg586 and Trp1250 whereas the catharanthine moiety displays a hydrogen-bond with Ser1097 (Figure 3c). Several van der Waals interactions provide further coordination of vinblastine within the TMD. The cryo-EM structure of P-glycoprotein (ABCB1/P-gp) has been reported with two vinblastine molecules, positioned deeper within the TMD interface at a depth corresponding to the junction between the inner and outer leaflets of the plasma membrane^14^ (Supplementary Figure 6). In rMrp2 the formyl group of vincristine likely contributes to (and may drive) the 2:1 stoichiometry by interlocking the two drug molecules. This is in contrast to MRP1 and P-gp wherethe vincristine formyl group does not make any intermolecular contacts with the second molecule and the presence of two vinblastine molecules in the P-gp structure is not mediated by the methyl group.

#### *Epipodophyllotoxin glycosides* - teniposide

We next examined teniposide, a semisynthetic glucoside belonging to the epipodophyllotoxin glycoside family of drugs. It is characterised by the epipodophyllotoxin core scaffold, a podophyllotoxin-derived structure modified with a glycosidic moiety. We resolved the cryo-EM structure of teniposide at 2.80 Å resolution (Supplementary Figure 7). In contrast to the vinca alkaloid complexes, density for the full transporter was well resolved including TMD0. Inspection of the coulomb maps revealed density for one teniposide molecule and CHS (Figure 4a). The teniposide spans both the drug binding sites 1 and 2 and the CHS is found in the H-pocket, similar to our previously reported probenecid bound structure^6^ (Figure 4b). The epipodophyllotoxin moiety is primarily stabilised by its lactone group through hydrogen bonds with Gln1246 and Trp1250 (Figure 4c). The *β*-D-glucopyranosyl ring is hydrogen bonded by Tyr433. The phenyl ring is hydrogen bonded by Arg586 and the two methoxy groups are stabilised by van der Waals interactions. The thiophene group that defines teniposide, displays a van der Waals interaction with Ser598. Extensive van der Waals interactions provide further coordination of the teniposide within the TMD (Figure 4c).

**Figure 4.**
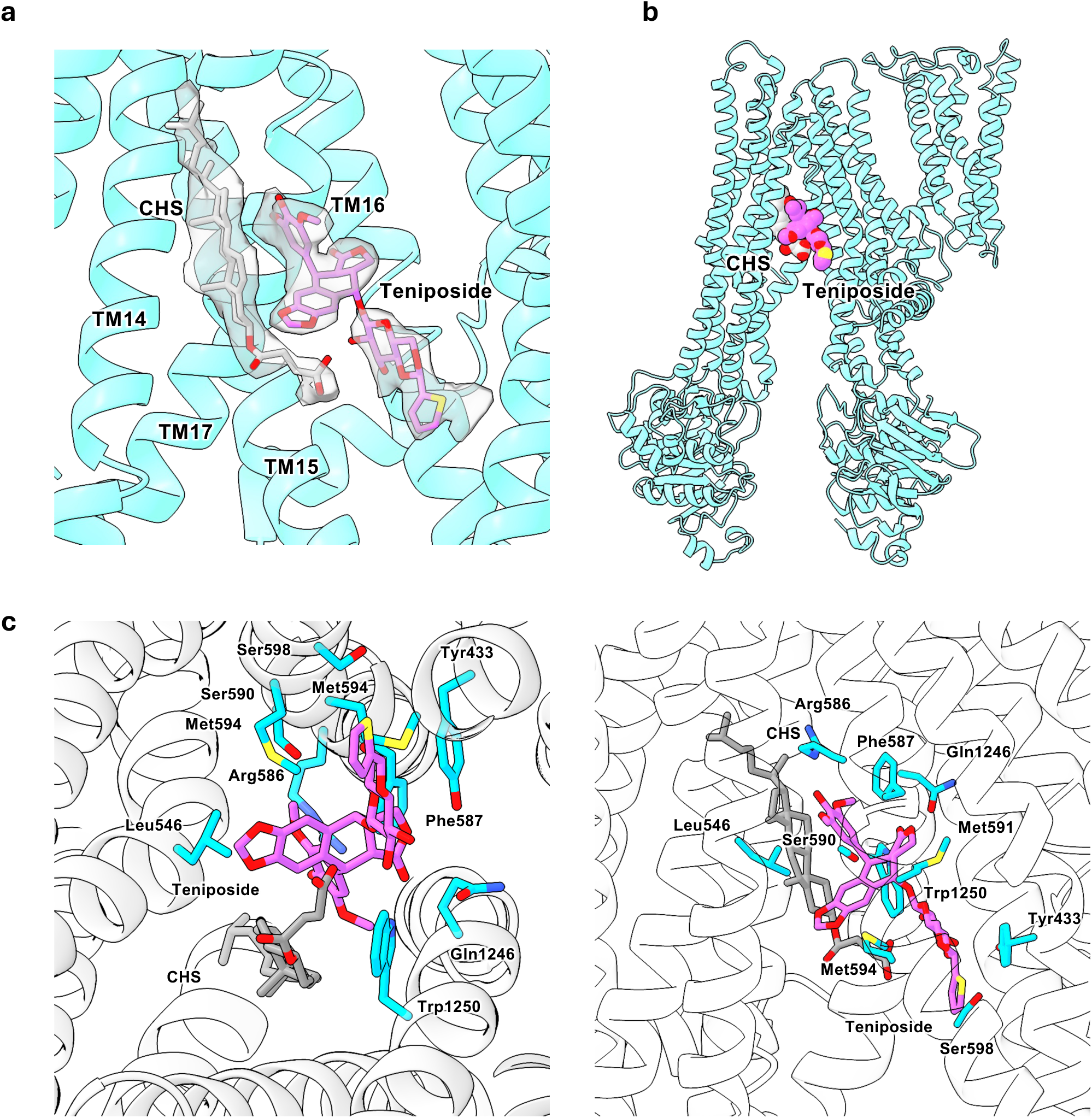
Structural basis of teniposide recognition by rMrp2. (a) Coulomb density for teniposide within the TMD, with the fitted ligand shown in pink and CHS in grey. (b) Overall structure of rMrp2 in the teniposide-bound state. (c) Close-up views of the binding pocket from two orientations, with coordinating residues shown as cyan sticks.

#### *Indene acetic acid class of NSAIDs* - sulindac

To extend the analysis beyond anticancer agents, we examined the prodrug sulindac, an NSAID of the arylalkanoic acid (indene acetic acid) class. Sulindac is extensively metabolised in the liver. The cryo-EM structure of rMrp2-sulindac was determined at 3.26 Å resolution (Supplementary Figure 8). The TMD contains two sulindac molecules (Figure 5a); one located at the interface between drug□binding pockets 1 and 2, and the other near the H□pocket (Figure 5b). The indene moiety is stabilised by van der Waals interactions between its fluorine atom and Tyr433 and Leu436, as well as *τ*-stacking with Phe378 (Figure 5c). The carboxylate of the indene moiety, which is essential for its NSAID activity, is hydrogen bonded by Arg1201. His328 and Gln443 hydrogen bond the prodrug moiety, phenyl-methylsulfinyl. The sulindac molecule near the H□pocket is stabilised by hydrogen bonds between Tyr433 and the phenyl-methylsulfinyl group, and His586 with the carboxylate of the indene moiety (Figure 5c). Both drugs display extensive van der Waals interactions with TMDs 1 and 2, and the two sulindac molecules engage in π□stacking interactions between their indene and phenyl□methylsulfinyl groups (Figure 5c).

**Figure 5.**
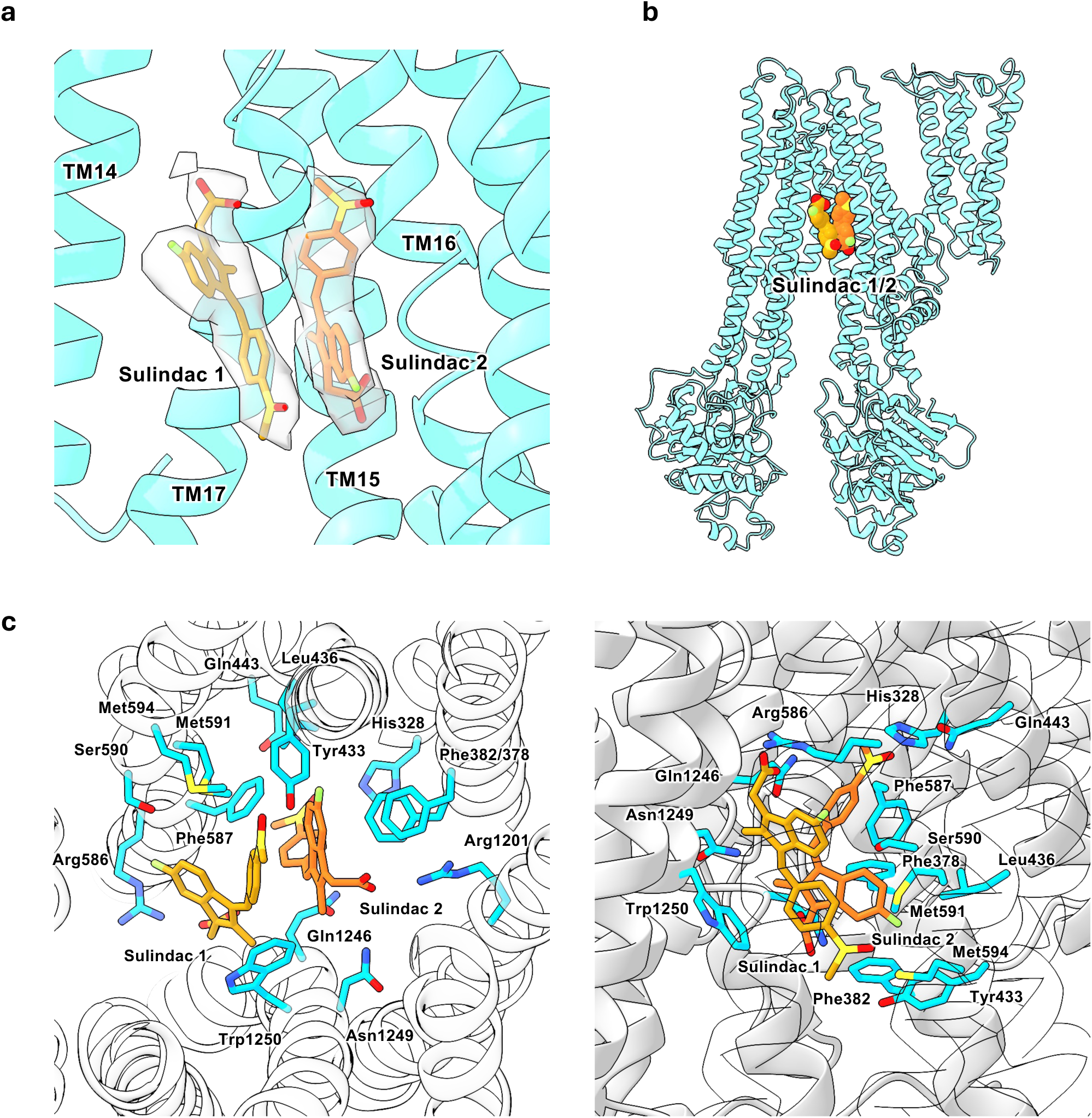
Structural basis of sulindac recognition by rMrp2. (a) Coulomb density (grey surface) and fitted models of two sulindac molecules bound within the TMD in different poses, shown in dark and light orange, respectively. (b) Overall structure of rMrp2 in the sulindac-bound state. (c) Close-up views of the binding pocket from two orientations, with coordinating residues shown as cyan sticks.

#### *Sulfonamide derivatives* - bosentan

To further test how rMrp2 accommodates chemically distinct non-oncology drugs, we next examined bosentan, a synthetic, non-peptide, arylsulfonamide molecule, characterised by a diaryl sulfonamide core bearing multiple alkyl□substituted phenyl groups. The cryo-EM structure of rMrp2-bosentan was determined at 3.41 Å resolution (Supplementary Figure 9). Most features of rMrp2 were resolved, except for the TMD0 which was disordered, similar to the vincristine- and vinblastine-bound structures. Inspection of the TMD, revealed coulomb density for three bosentan molecules (Figure 6a). One bosentan molecule is positioned near the H□pocket and it is stabilised by polar interactions involving Gln539 and the diaryl sulfonamide core, and extensive van der Waals interactions (Figure 6b and c). The second molecule sits within drug□binding pocket 2 and it is stabilised by two salt-bridges between its diaryl sulfonamide group and Arg1145 and Arg1253, respectively, a *τ*-stacking interaction with Phe382 and its alkyl□substituted phenyl group; extensive van der Waals interactions with the TMD provide further interactions (Figure 6b and c). The third molecule is coordinated by a hydrogen bond with Arg1253 and van der Waals interactions with TMDs 1 and 2. This third molecule is localised to the interface between the two other bosentan molecules. The three bosentan molecules also display extensive intermolecular *π*-stacking interactions between their diaryl sulfonamide and alkyl□substituted phenyl groups (Figure 6b and c).

**Figure 6.**
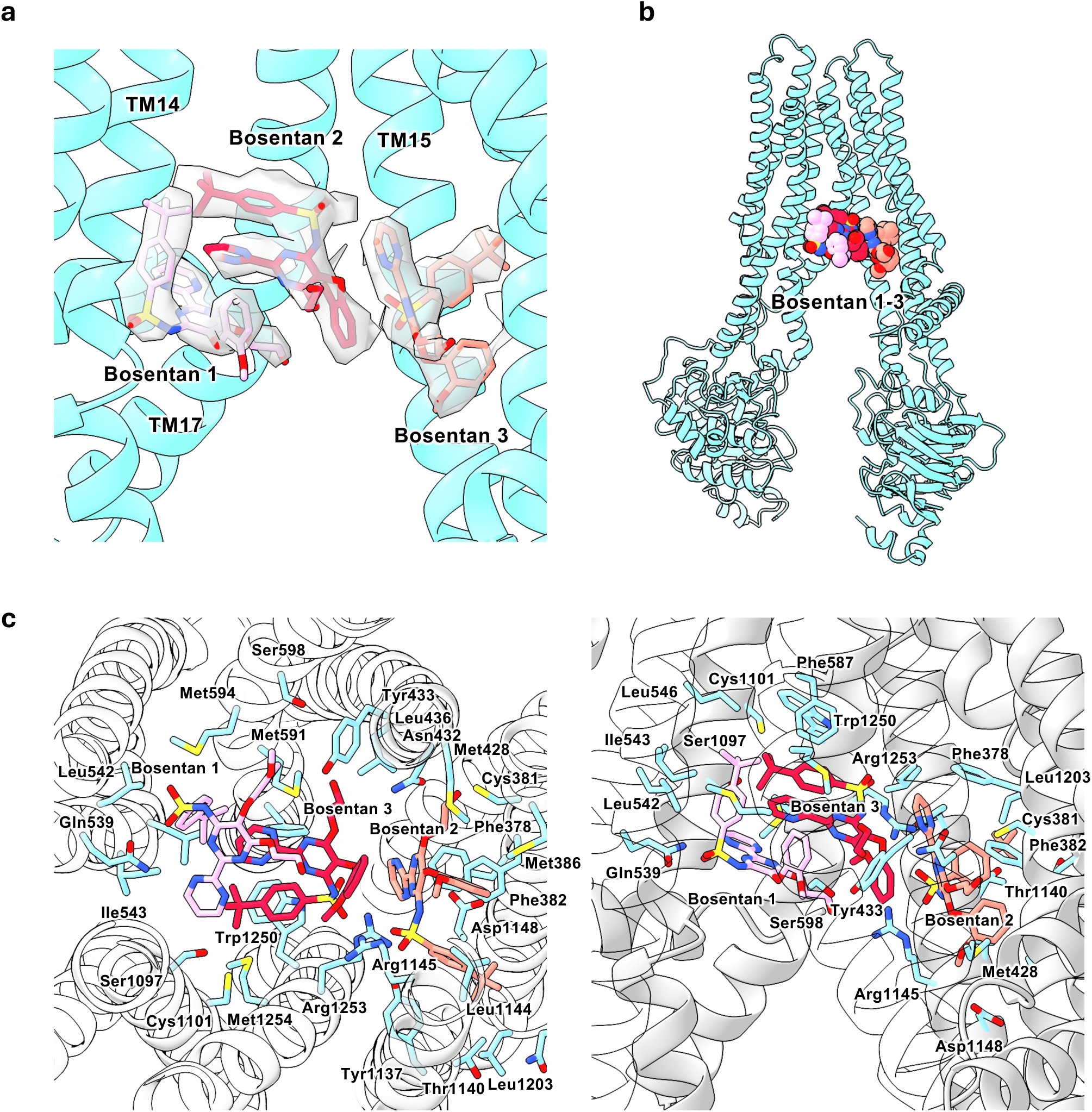
Structural basis of bosentan recognition by rMrp2. (a) Coulomb density (grey surface) and fitted models of three bosentan molecules bound within the TMD, shown in pink, red and salmon sticks, respectively. (b) Overall structure of rMrp2 in complex with bosentan. (c) Close-up views of the binding pocket, highlighting drug-drug interactions among the bound bosentan molecules; coordinating residues are shown as cyan sticks.

#### *Heteroaryl□sulfonamides* – MK-571

We also established the binding mode of the non-specific ABCC/MRP inhibitor MK-571 (https://transportal.compbio.ucsf.edu/compounds/mk-571/). MK-571 is a synthetic quinoline derivative containing a 4-oxo-quinoline core with a 3-substituted carboxylic acid and an aryl-sulfonamide side chain. It has been characterised as an MRP1 (and related ABCC/MRP) inhibitor, blocking leukotriene C4 export and other organic anion efflux, and is widely used as a tool compound to study multidrug resistance and leukotriene-dependent signalling ^15^. The cryo-EM structure of the rMrp2 bound to MK-571 was determined at 3.02 Å resolution (Supplementary Figure 10). We resolved coulomb density for one MK-571 molecule and CHS (Figure 7a). MK-571 is bound within the H-pocket (Figure 7b), where the chloroquinoline moiety is stabilised by hydrogen-bonding interactions with the backbone of Asn1243 and side chain of Thr1247, while its chlorine and pyridine ring engage in polar interactions with Trp1250 (Figure 7c). Extensive van der Waals interactions provide further stabilisation. Comparison with the leukotriene C4 bound structure shows significant overlap with MK-571 and its inhibitory action is very likely from direct competition for the same binding site (Supplementary Figure 11).

**Figure 7.**
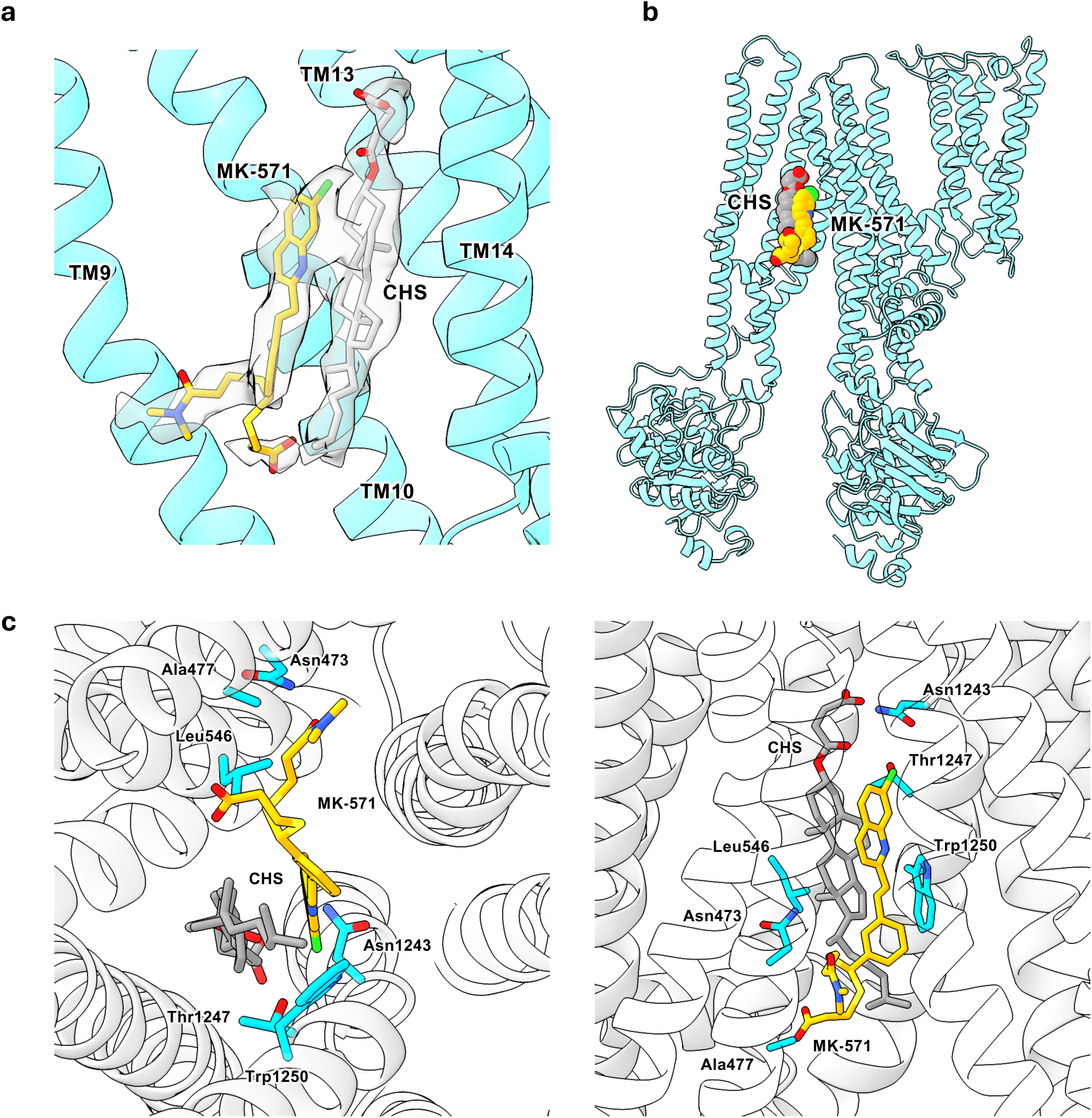
Structural basis of MK-571 recognition by rMrp2. (a) Coulomb density for MK-571 within the TMD, with the fitted ligand shown in yellow and CHS in grey. (b) Overall structure of rMrp2 in the MK-571-bound state. (c) Close-up views of the binding pocket, with coordinating residues shown as cyan sticks.

**Figure 8.**
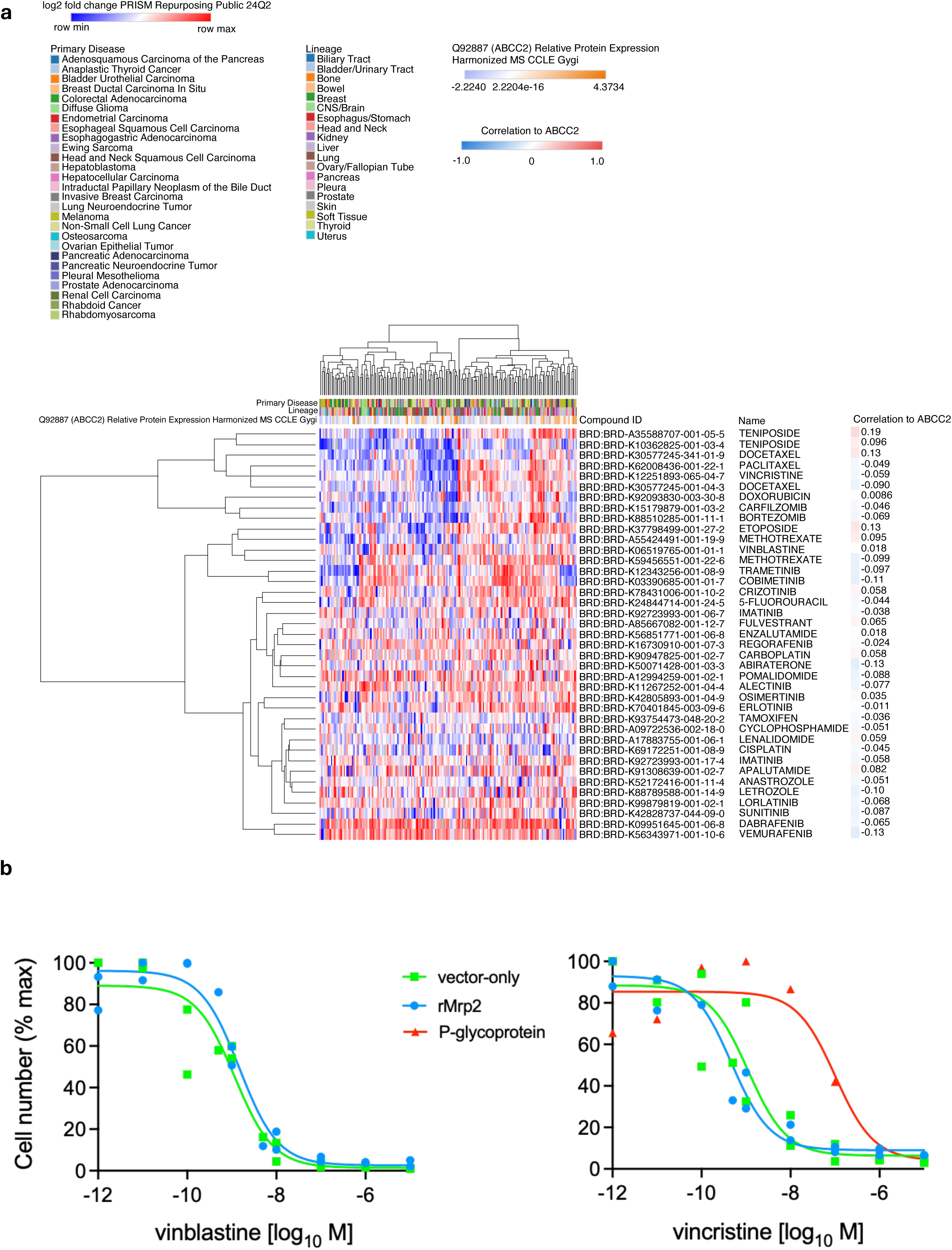
Lack of correlation between ABCC2 protein abundance and cancer drug response across DepMap cell lines. (a) Heatmap showing log2 fold-change drug response values from the PRISM Repurposing Public 24Q2 dataset across cancer cell lines, ordered by hierarchical clustering. Rows represent individual compounds and columns represent cancer cell lines. Cell lines are annotated by primary disease and lineage (top color bars). ABCC2 (Q92887) relative protein expression, from harmonized MS-CCLE proteomics data, is shown as an annotation bar above the heatmap. For each drug, the Pearson correlation coefficient between drug response and ABCC2 protein abundance across cell lines is shown on the right. Overall, correlations between ABCC2 protein expression and drug response are weak and not statistically significant, indicating no strong association between ABCC2 abundance and sensitivity to the analysed cancer drugs. (b) Stable rMrp2 expression in HEK293-FlpIn cells does not confer resistance to vinblastine or vincristine. Nonlinear regression analysis of cell number after three-day culture in the presence of increasing concentrations of vinblastine (left hand graph) or vincristine (right hand graph). Data are normalised to maximum cell number. The IC_50_ for vinblastine calculated for the rMrp2 expressing cells (1.48 nM 95% CI [0.93, 2.4] n=2) is statistically indistinguishable (extra sum-of-squares F test) from the vector only cells (1.16 nM 95% CI [0.45, 2.8] n=2). The IC_50_ for vincristine killing of HEK293-FlpIn-rMrp2 (0.47 nM 95% CI [0.24, 0.92] n=2) and HEK293-FlpIn-vector cells (1.11 nM 95% CI [0.30, 4.7] n=2) are also statistically indistinguishable. For comparison, vincristine resistance conferred by P-gp expression in HEK293-FlpIn-ABCB1 cells^24^, described by an IC_50_ for vincristine of 101.1 nM 95% CI [13.2, 884] n=1) is significantly different (p < 0.0001) to both the control and rMrp2 expressing cell lines.

### MRP2 abundance does not correlate with drug response in cancer cell lines

To investigate whether MRP2 is associated with drug response in cancer models, we integrated DepMap PRISM (24Q2) drug-response profiles with harmonized MS-CCLE proteomics^16^ and found that MRP2 protein abundance exhibits uniformly weak, non-significant correlations with responses to a broad panel of oncology agents, including vincristine, vinblastine and teniposide, across a combined set of diverse lineages (maximum |r|≈0.19) (Figure 9a), arguing against a simple association between MRP2 expression and cytotoxic drug sensitivity *in vitro*. We tested this further by generating a cell line that expresses a single copy of rMrp2 under control of the CMV promoter and measuring its response to cytotoxic challenge. Rat Mrp2 did not confer a measurable resistance phenotype to the vinca alkaloids in our cell toxicity/resistance assays, while P-gp, expressed from the same locus, conferred 100-fold resistance to vincristine compared to the vector-only control (Figure 9b). These empirical data agree with the DepMap data, despite our structural and functional data showing that rMrp2 interacts with, and is inhibited by, the vinca alkaloids. Together, these findings are consistent with the primary impact of MRP2 being pharmacokinetic and subject to drug-drug interactions, rather than a robust cancer cell-intrinsic resistance mechanism, suggesting that any MRP2-mediated chemoresistance is likely context-dependent.

**Figure 9.**
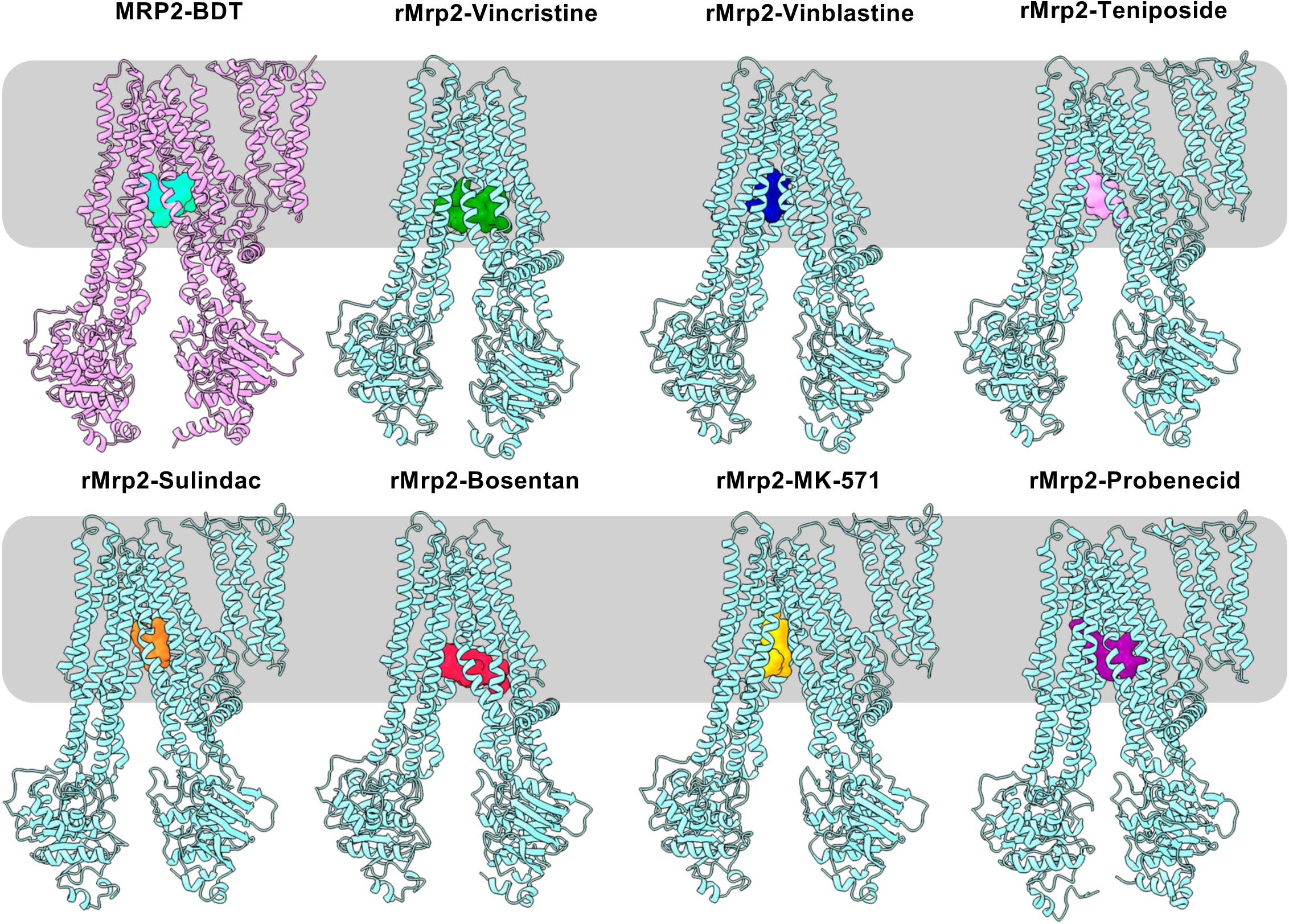
Comparison of ligand binding poses of rMRP2 with MRP2. Cryo-EM structures of rMrp2 bound to vincristine (green), vinblastine (blue), teniposide (pink), sulindac (orange), bosentan (crimson), MK-571 (yellow) and probenecid (PDB: 8RQ4, purple) are shown with rMrp2 depicted as aquamarine cartoon. Lipids are shown in grey. Each ligand is shown as a coloured surface. The structure of MRP2 bound to bilirubin ditaurate (BDT) (PDB: 8JQX) is shown in the top left; MRP2 is shown as violet cartoon and BDT as cyan surface. All structures are aligned on the transmembrane domain and displayed in the same orientation, with the approximate position of the lipid bilayer indicated by the grey box.

## Discussion

In this study, we investigated the structural basis of ligand recognition by rMrp2 using a panel of clinically used drugs from diverse chemical classes. The high degree of sequence conservation of the transmembrane drug-binding cavity between rat and human orthologues (98% identity) indicates that the recognition principles identified here are likely to be relevant to human MRP2. As shown by our functional data, rMrp2 recognises the same drugs as the human protein supporting its use as a suitable model for structural analysis.

A central finding of this work is that rMrp2 polyspecificity cannot be readily explained by a shared pharmacophore or a simple set of conserved ligand contacts. The compounds analysed here are chemically diverse, with pairwise Tanimoto coefficients below 10% for all comparisons except vincristine and vinblastine, which are highly similar at 88% (Supplementary Figure 12). Maximum common substructure analysis identified a tolyl moiety, toluene, as the only recurrent feature across the broader ligand set (Supplementary Figure 12), but this functional group is unlikely to drive specificity because its position, local coordination and binding pose differ markedly between structures. Instead, the cryo-EM structures reveal a recognition mechanism in which chemically distinct ligands converge on overlapping regions of the transmembrane cavity while engaging the protein through different poses, contact networks and stoichiometries (Figure 9 and Supplementary Figure 13).

The vinca alkaloids illustrate this principle particularly clearly. Despite their close structural similarity, vincristine and vinblastine adopt distinct binding poses and stoichiometries in rMrp2. Previous work on MRP1 resolved in complex with two vincristine molecules was interpreted as non-physiological^13^. In contrast, MRP4 has been reported to bind only a single vincristine molecule^17^, indicating that this behaviour is not universal across the ABCC family. In our structure, the two vincristine molecules occupy sites within the rMrp2 transmembrane cavity distinct from those observed in MRP1 and 4. We propose that it is unlikely to be coincidental and may point to a complex relationship between vinca alkaloids and their recognition by ABC transporters. The differences in the physicochemical properties of the drugs do not appear to contribute to the differences in stoichiometry as both vincristine and vinblastine can exist either in a 1:1 or 2:1 drug to ABC transporter ratio. More broadly, our structures show that ligand stoichiometry and binding geometry vary substantially across compounds (Figure 9 and Supplementary Figure 13). Bosentan and probenecid both contain a benzenesulfonamide pharmacophore (Supplementary Figure 12), yet they bind rMrp2 in strikingly different ways, with bosentan adopting a 3:1 stoichiometry and probenecid a 2:1:1 stoichiometry in the presence of cholesteryl hemisuccinate. These observations suggest that both ligand stoichiometry and binding geometry may influence transport efficiency, and that these relationships likely vary across ABC transporters and among individual substrates. Importantly, while these compounds stimulate rMRP2’s ATPase activity, the magnitude of this stimulation does not correlate with ligand stoichiometry, or the presence of CHS, indicating that catalytic activation occurs independently of binding pocket occupancy and architecture.

By mapping interactions from all rMrp2 drug-bound structures onto the transporter, we can delineate a pliant drug-binding cavity and identify residues that contribute to ligand recognition and stabilisation, thereby providing a structural basis for polyspecific substrate binding by the TMDs (Figure 10 and Supplementary Figure 14). Rather than occupying discrete sites, the drugs bind within a shared pocket of approximately 4,700A^3^. Phe587 and Trp1250 make the most recurrent contacts, interacting with 5 of 7 and 7 of 7 drugs, respectively (Figure 10).

**Figure 10.**
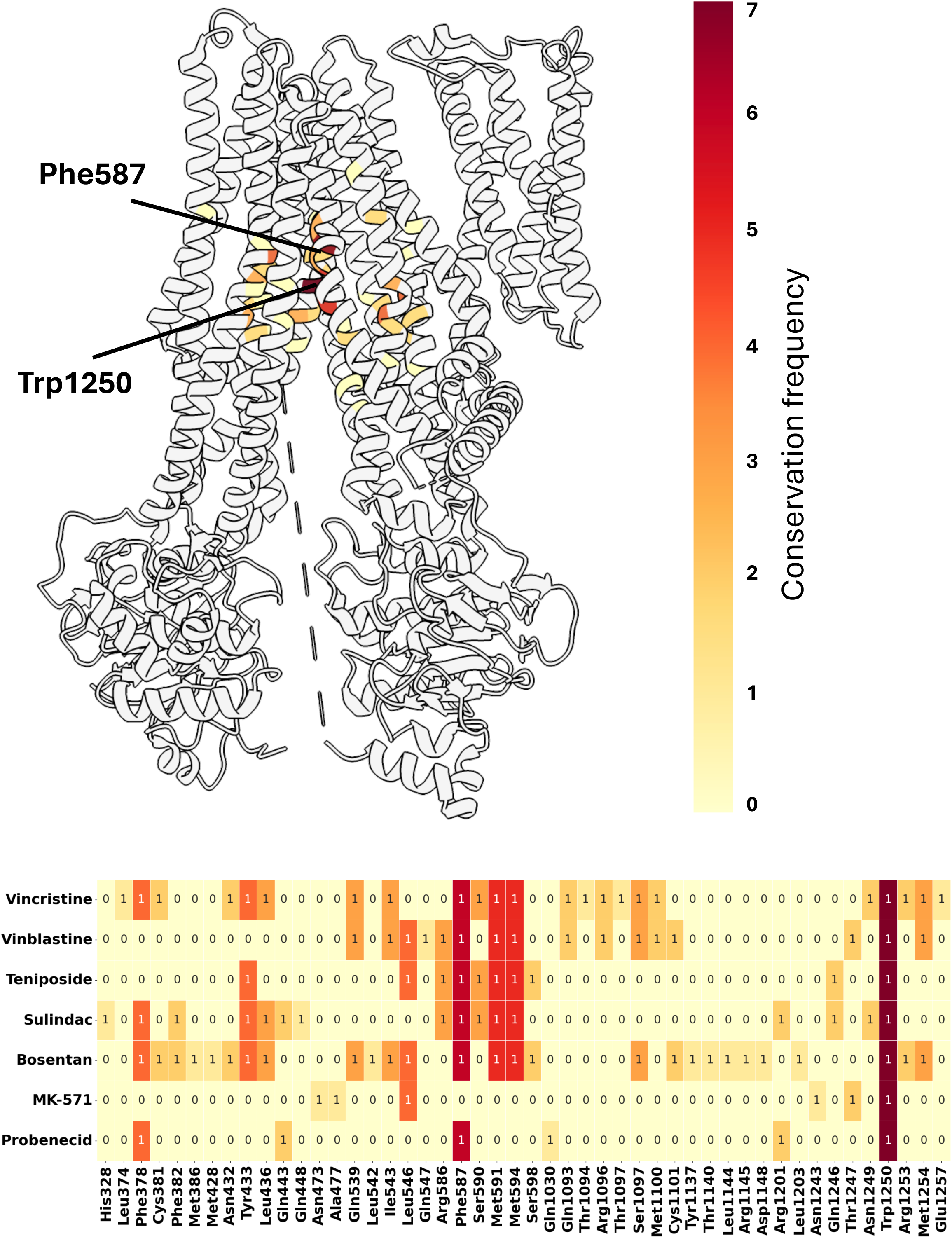
Heatmap of residue-ligand interactions and conservation. The heatmap summarises the binding profiles of all drug-bound structures across identified amino acid residues. Colour intensity, ranging from light yellow to dark red, reflects conservation frequency. Interacting residues are mapped onto the rMRP2 structure, with Phe587 and Trp1250 indicated. Together, these residues define the boundaries of the drug-binding region within the TMD.

Previous mutagenesis of the equivalent residues in MRP2, Phe591 and Trp1254, caused only modest reductions in bilirubin ditaurate-stimulated ATPase activity^18^, suggesting that no single residue is solely responsible for substrate recognition. By contrast, mutation of the equivalent tryptophan in MRP1, Trp1246, abolished transport activity in HeLa cells^19^. Notably, Trp1250 in rMrp2, and the corresponding residue in MRP2 and MRP1, forms a hydrogen bond with Ser1097 in the R-domain in the autoinhibited state^6^, indicating a broader role in transporter regulation^18, 6^.

Superposition of the apo autoinhibited and drug-bound structures shows that Trp1250 swings into the cavity region otherwise occupied by the R-domain (Supplementary Figure 15). Its conformation is conserved across the drug-bound states, except in the bosentan-bound structure, where one of the three bound molecules displaces it further outward. Rotamer analysis using the Dunbrack library indicates that the autoinhibited conformation corresponds to a low-prevalence rotamer of approximately 1.5% probability, whereas the drug-bound conformation is much more common at approximately 30%. This suggests that the autoinhibited geometry is locally stabilised despite being intrinsically less favoured. Consistent with our model in which R-domain disengagement relieves autoinhibition and permits substrate binding, loss of transport activity upon mutation of the equivalent residue in MRP1, Trp1246, may therefore reflect perturbation of the autoinhibited state rather than loss of a direct ligand-contacting side chain. In this view, the conserved tryptophan defines the innermost boundary of the cavity exposed upon relief of autoinhibition, and its recurrent contact with chemically diverse ligands likely reflects cavity geometry rather than a specific role in substrate recognition.

Taken together, these observations support a model in which MRP2 polyspecificity arises primarily from a predominantly hydrophobic transmembrane cavity that can accommodate ligands of different sizes, shapes and topology without requiring a common pharmacophore. Across the structures, most ligand contacts are mediated by van der Waals interactions, and overlapping recognition appears to depend more on general hydrophobic complementarity than on a conserved network of polar interactions. This model is also consistent with the broader substrate repertoire of MRP2, which includes structurally unrelated compounds such as methotrexate^3^, saquinavir^20^ and SN-38 glucuronide^21^. We therefore propose that MRP2 recognises chemically heterogeneous substrates through a permissive cavity which is further modulated by ligand size, shape and local charge distribution.

Our ATPase measurements and structures further suggest that several parent drugs can be directly recognised by rMrp2 and may therefore be substrates as well as inhibitors. However, evidence of direct transporter engagement did not translate into a measurable resistance phenotype in our cellular assays. Stable expression of rMrp2 in HEK293 cells failed to confer resistance to vincristine or vinblastine, despite clear evidence of binding in structural experiments, stimulation of the ATPase *in vitro* and inhibition of the transport of CDF *in cellulo*. One interpretation is that these compounds are recognised but transported too slowly, or with insufficient capacity, to overcome their intrinsic cytotoxicity. Another possibility is that incomplete activation of Mrp2 through phosphorylation-dependent regulation limits effective efflux under our experimental conditions (despite measurable efflux of CDF). Regardless of mechanism, our data indicate that ligand binding alone is not sufficient to infer robust transporter-mediated chemoresistance.

This conclusion is consistent with our analysis of publicly available drug-response datasets from the DepMap portal, which did not reveal a strong association between ABCC2 abundance and sensitivity to a panel of 27 anticancer agents across diverse cancer cell line models. Although MRP2 has often been discussed in the context of chemotherapy resistance, the clinical evidence remains limited, and much of the existing support derives from overexpression systems^22,23^. Our results argue for a more cautious interpretation: cytotoxic drugs can engage and inhibit MRP2, but such interactions do not necessarily generate a detectable resistance phenotype.

The physiological implications of our findings may therefore lie less in multidrug resistance than in hepatobiliary xenobiotic handling. Given its canalicular localisation, MRP2 is well positioned to contribute to hepatic protection by exporting drugs, metabolites and conjugates into bile, in parallel with other efflux systems such as P-gp. In this context, overlapping ligand recognition within the transmembrane cavity provides a plausible mechanistic basis for transporter-mediated drug-drug interactions. Competition between co-administered compounds could reduce the efflux of lower-affinity substrates and increase intracellular exposure, particularly under polypharmacy conditions. Such effects may be relevant to altered hepatobiliary clearance and, in some settings, to susceptibility to cholestatic or drug-induced liver injury, although this was not directly examined here. The consequences of these interactions are likely to be further shaped by the phosphorylation state of MRP2 and by coordination with other canalicular transporters.

Overall, our results define a structural basis for MRP2 polyspecificity and show that it recognises chemically diverse ligands through a predominantly hydrophobic transmembrane cavity capable of supporting multiple binding poses and stoichiometries. These findings provide a mechanistic framework for understanding how MRP2 engages chemically diverse ligands and underscores the need to distinguish transporter binding from efficient transport when interpreting the pharmacological consequences of ABC transporter–drug interactions.

## Acknowledgments

The cryo-EM experiments were performed at the UK National Electron Bio-Imaging Centre (eBIC) under visit NT29493. We would like to acknowledge the Research Complex at Harwell for access. This project made use the Computing Platform for Electron Microscopy at Imperial College funded by the BBSRC Mid-range equipment Initiative 22ALERT BB/X019284/1 to KB.

## Author contributions

KB and KJL conceived and supervised the study. NI purified the rMrp2, measured the ATPase activity, determined the cryo-EM structure, performed model building and refinement. KB analysed the structures and performed pharmacophore analysis. HK, EJM, LY and KJL engineered rMrp2 intergration into HEK293 cells, performed competition and cell survival assays. TIR performed bioinformatics analysis. KB and KJL wrote the manuscript with input from all authors.

## Data availability

The cryo-EM maps have been deposited in the Electron Microscopy Data Bank (EMDB) under accession codes EMD-80514 (vincristine), EMD-80515 (vinblastine), EMD-80516 (teniposide), EMD-80517 (sulindac), EMD-80518 (bosentan) and EMD-80519 (bosentan). The atomic coordinates have been deposited in the Protein Data Bank (PDB) under accession codes 26BN (vincristine), 26BO (vinblastine), 26BP (teniposide), 26BQ (sulindac), 26BR (bosentan) and 26BS (bosentan). Raw movies for vincristine, vinblastine, teniposide, sulindac, bosentan and MK-571 were submitted to Electron Microscopy Public Image Archive (https://www.ebi.ac.uk/pdbe/emdb/empiar/) with IDs EMPIAR-1xx, EMPIAR-2xx, EMPIAR-3xx, EMPIAR-4xx, EMPIAR-5xx and EMPIAR-6xx, respectively.

## Materials and Methods

### Generation of the HEK293-FlpIn-Mrp2 and control cell lines

Plasmid cDNA5/FRT-Mrp2 was engineered by a modified Gibson reaction from three fragments amplified with Q5 High Fidelity polymerase (New England Biolabs (NEB)). The 5’ end of Mrp2 cDNA was amplified from pwaldo-rMrp2-GFP^6^ using the rMrp2 forward and rMrp2 2311R primers (all primers are described in Supplementary Table 3), while the 3’ end of the cDNA was amplified using the Mrp2 reverse primer (modified to re-introduce a stop codon which had been deleted when the cDNA was fused to the coding sequence for GFP in pwaldo-rMrp2-GFP) and Mrp2 2330F primer. Plasmid cDNA5-FRT (ThermoFisher Scientific) was amplified with forward and reverse primers. The pcDNA5-FRT forward primer was also adapted to include the stop codon for rMrp2. PCR products were separated by agarose gel electrophoresis, recovered from the gel and assembled using NEBuilder HiFi DNA Assembly (NEB).

HEK293-FlpIn cells with a single genomic FLP recombinase integration site were purchased from ThermoFisher Scientific. The cells were maintained in growth medium, Dulbecco’s modified Eagle medium (DMEM) high glucose (ThermoFisher Scientific) supplemented with 10% foetal bovine serum (FBS; Sigma-Aldrich) and 100 units/mL of penicillin and 100 µg/mL of streptomycin (ThermoFisher Scientific). The cells were co-transfected with pcDNA5-FRT-rMrp2 and pOGG44 (ThermoFisher Scientific) encoding the FLP recombinase. After 48hrs, stable recombinants were selected with 150 *μ*g/ml hygromycin B (TOKU-E) and once established, maintained with 75 *μ*g/ml hygromycin B. The hygromycin resistant pcDNA5/FRT vector-only cell line was generated in the same manner.

The respective negative and positive controls HEK293-FlpIn-vector and HEK293-FlpIn-ABCB1 were generated similarly and described previously^24^.

### Western Blot analysis

Cultured HEK293 FlpIn cells were harvested and solubilised in lysis buffer (Tris pH6.8 100mM, EDTA 10mM, SDS 2%, Triton X100 0.1%, Complete protease inhibitors (2x; Sigma-Aldrich). Proteins were separated in a 7.5% polyacrylamide gel, transferred onto PVDF, blocked in 5% powdered milk solution and incubated with primary antibody (mouse monoclonal anti-rMrp2, M2 III-6 (Abcam)).

### *In cellulo* rMrp2 efflux and inhibition assay

HEK293-FlpIn-rMrp2 (or vector-only) cells were harvested with 0.5 ml of TrypLE express enzyme (ThermoFisher Scientific) and quenched with 4.5 ml of growth medium (without antibiotics). The cells (1 x 10^6^ cells/ml) were loaded with 20 µM 5(6)-carboxy-2′,7′-dichlorofluorescein diacetate (CDFDA; Sigma-Aldrich) for 10 min at 37°C. Cells were pelleted by centrifugation (180 × g for 1 min) and resuspended at 1.3 x 10^6^ cells/ml in growth medium without antibiotics. An aliquot (75 µL) was immediately placed on ice as the 5(6)-carboxy-2′,7′-dichlorofluorescein (CDF) loaded control. Remaining aliquots were mixed with 75 µL of growth medium (without antibiotics), with or without potential inhibitor at a range of empirically determined concentrations. After a 15-minute incubation at 37 °C, cells were pelleted as before and resuspended in 300 µL of cold growth medium (without antibiotics) and placed on ice until flow cytometry (ACEA NovoCyte flow cytometer; Agilent Technologies). Analysis was in FlowJo (MacOS Version 10.8.2; Becton Dickinson) where size, granularity and singlets were gated on an untreated cell sample and applied to all samples.

CDF florescence was recorded in the green fluorescence (FITC) channel. For each condition, median fluorescence values were subtracted from the loaded control (zero efflux) and calculated as a percentage of the no-drug control (max efflux). Percentage rMrp2-activity values were modelled by non-linear regression (three-parameter curve fit in GraphPad PRISM; MacOS Version 10.4.0) with the baseline constrained to 0, to determine the IC_50_ values for each drug. Efficacy was calculated as the lowest activity of rMrp2 in the presence of drug compared to the top of the curve. Triplicate biological repeats were completed for each compound.

### *In cellulo* drug resistance assay

Cells were seeded (10,000 cells per well) into a 96-well dish in 50 *μ*L growth medium (as described above but without the antibiotics). Vincristine or vinblastine was added in a 50 *μ*l volume to achieve a final concentration ranging from 0 to 10 *μ*M. The cells were cultured for a further 72 h, after which the medium was aspirated, the cells were detached with 30 *μ*L TrypLE Express, before quenching with 75 *μ*L growth medium (with only 1% FBS and no antibiotics) for flow cytometry. Cells of normal size and granularity were gated in the zero-drug condition. This gate was applied to all samples for counting in an ACEA NovoCyte flow cytometer (Agilent Technologies, Santa Clara, CA, USA). Cell number data were analysed in Graphpad PRISM (MacOS version 10.4.0). Three parameter curve fitting was unconstrained and statistical analysis of the IC_50_ was by extra sum-of-squares F test.

### Expression and purification of rMrp2

Rat Mrp2 (rMrp2) was expressed in *Saccharomyces cerevisiae* from pwaldo-rMrp2-GFP and purified as previously described^6^. In brief, isolated membranes were extracted in 50 mM Tris-HCl pH 8.0, 300 mM NaCl, 2 mM MgCl□, 2 mM DTT, 20% glycerol supplemented with 2% LMNG and 0.4% CHS for 90 min at 4 °C. The protein was purified using a His-Trap column and the detergent was exchanged using a Superose 6 10/300 column (50 mM Tris-HCl pH 8.0, 150 mM KCl, 2 mM MgCl□, 2 mM DTT and 0.06% GDN). Peak fractions were collected and concentrated to 3-4 mg/ml.

### ATPase activity measurement of rMrp2 with drugs

The ATPase activity of rMrp2 was measured using the EnzChek Phosphate Assay Kit (Thermo Fisher Scientific), following our previously described protocol^6^. In brief, rMrp2 was reconstituted into destabilised liposomes consisting of bovine liver total lipid extract (Avanti Polar Lipids), and the linear ATP hydrolysis rate over the first 1,200 s was measured in the apo state or in the presence of drugs. Reactions were carried out in the presence of 5 mM ATP and 5 mM MgCl□ at room temperature. For ligand-induced ATPase activity assays, drugs were dissolved in 100% DMSO to prepare 100 mM stock solutions and added to the reactions at the following final concentrations: 0.1 mM for MK-571, sulindac, and teniposide, and 1 mM for probenecid, vinblastine, vincristine and bosentan.

### Grid preparation and CryoEM data collection and processing

3.5 µL of the purified rMrp2 at 3-4 mg/mL was applied to a glow-discharged Quantifoil R1.2/1.3 300 mesh Cu Holey carbon grid treated with 3□mM fluorinated Fos-Choline-8 (Anatrace). The grid was blotted with a blotting force of 0 for 3 sec at 4°C and 95% humidity, and flash-frozen in liquid ethane using a Vitrobot Mark IV (Thermo Fisher Scientific). The frozen grid was stored in liquid nitrogen until data collection. CryoEM data were collected on 300 kV Titan Krios (Thermo Fisher Scientific) with a Falcon 4i detector and Selectris X energy filter or Gatan K3 camera at the Electron Bioimaging Center (eBIC), UK. The data acquisition parameters are shown in Supplementary Table 4.

### CryoEM Data Processing

CryoEM data were processed using cryoSPARC v4.6.1^25^. Each dataset was processed independently following the same general workflow. Movies were corrected for beam-induced motion using Patch Motion Correction, and CTF parameters were estimated using Patch CTF Estimation^25^. Initial particle picking was performed with Blob Picker on 500–1,000 micrographs without templates, and particles were extracted with binning state (pixel size: 1.5 – 3.5 Å/px). Extracted particles were subjected to 2D classification, and selected suitable 2D classes were used for Ab-Initio Reconstruction in 3 classes. A second round of 2D classification was performed on particles from the best ab-initio class, and the suitable 2D classes were used as templates for Template Picker. Template-picked particles were extracted with binning state and performed two rounds of Heterogeneous Refinement. Particles from the best 3D class were re-extracted to high-resolution pixel size (0.99 Å/px or 1.33 Å/px) and subjected to an additional round of Heterogeneous Refinement. The resulting class was further processed by 3D Classification into 4–10 classes. For the vinblastine and MK-571 datasets, 3D Classification was performed with a focused mask covering TMD1–2 to resolve ligand-bound states. The 3D volumes obtained from 3D Classification were used as references for Heterogeneous Refinement. Classes in which ligand density was observed were selected and subjected to Non-Uniform Refinement^26^. For the Vincristine, bosentan, and sulindac datasets, this map was taken as the final reconstruction. For the teniposide, vinblastine, and MK-571 datasets, Local CTF Refinement was subsequently performed. The teniposide dataset was additionally processed with Reference-Based Motion Correction. A second round of Non-Uniform Refinement was then performed to calculate the final maps. The resolution of each final map was estimated using Fourier Shell Correlation with a cutoff of 0.143, following the gold-standard criterion. Model building and refinement were performed starting from the rMrp2-probenecid bound structure (PDB ID: 8RQ4). The structure was fitted by rigid-body into the map using UCSF ChimeraX^27^ and real-space refined in phenix^28^. The models were then subjected to iterative cycles of refinement and manual rebuilding in COOT^29^. The structural models were analysed and validated using MolProbity^30^. The data processing and refinement statistics are shown in Supplementary Table 4.

### Drug response bioinformatic analysis

Drug response data were obtained from the DepMap1 PRISM Repurposing Public 24Q2 release and expressed as log2 fold-change values across cancer cell lines (https://depmap.org). ABCC2 (Q92887) protein abundance data were retrieved from the DepMap harmonized MS-CCLE proteomics dataset. Cell lines with available measurements for both drug response and ABCC2 protein expression were included in the analysis. For each compound, Pearson correlation coefficients were calculated between drug response values and ABCC2 protein abundance across cell lines. No additional stratification by cancer type was applied. Hierarchical clustering of drugs and cell lines was performed based on drug response profiles to visualize global patterns. Visualization was performed in the Phantasus tool2^31^.

