## Supplementary material for "Structural basis of polyspecific drug recognition by MRP2"

**Supplementary Text**

**HEK293-FlpIn-rMrp2 cells express rMrp2 which can efflux CDF**

Western blot analysis of whole cell lysates prepared from the recombinant HEK293-FlpIn-rMrp2 cells showed the presence of a high molecular weight protein species detected by anti-rMrp2 antibody (Supplementary Figure S3). The rMrp2 which is expressed from a single copy of the cDNA integrated into the genome of the FlpIn cells is also functional. CDFDA is a hydrophobic non-fluorescent compound that can cross the plasma membrane. In the cytosol the diacetate moiety is hydrolysed by esterases to release fluorescent CDF which is a transport substrate of rMrp2. The HEK293-FlpIn-rMrp2 accumulate less CDF than the vector-only cells because rMrp2 is also actively effluxing the CDF as it is made during the CDFDA loading step (Supplementary Figure S3). However, the loaded cells continue to efflux CDF if incubated at 37$^{o}$C (compare the histograms for the 'loaded' and 'no drug' samples in Figure 1c). The efflux activity, post loading, can be inhibited by incubation in the presence of increasing concentrations of drug (Figure 1c/d) to estimate potency and efficacy.

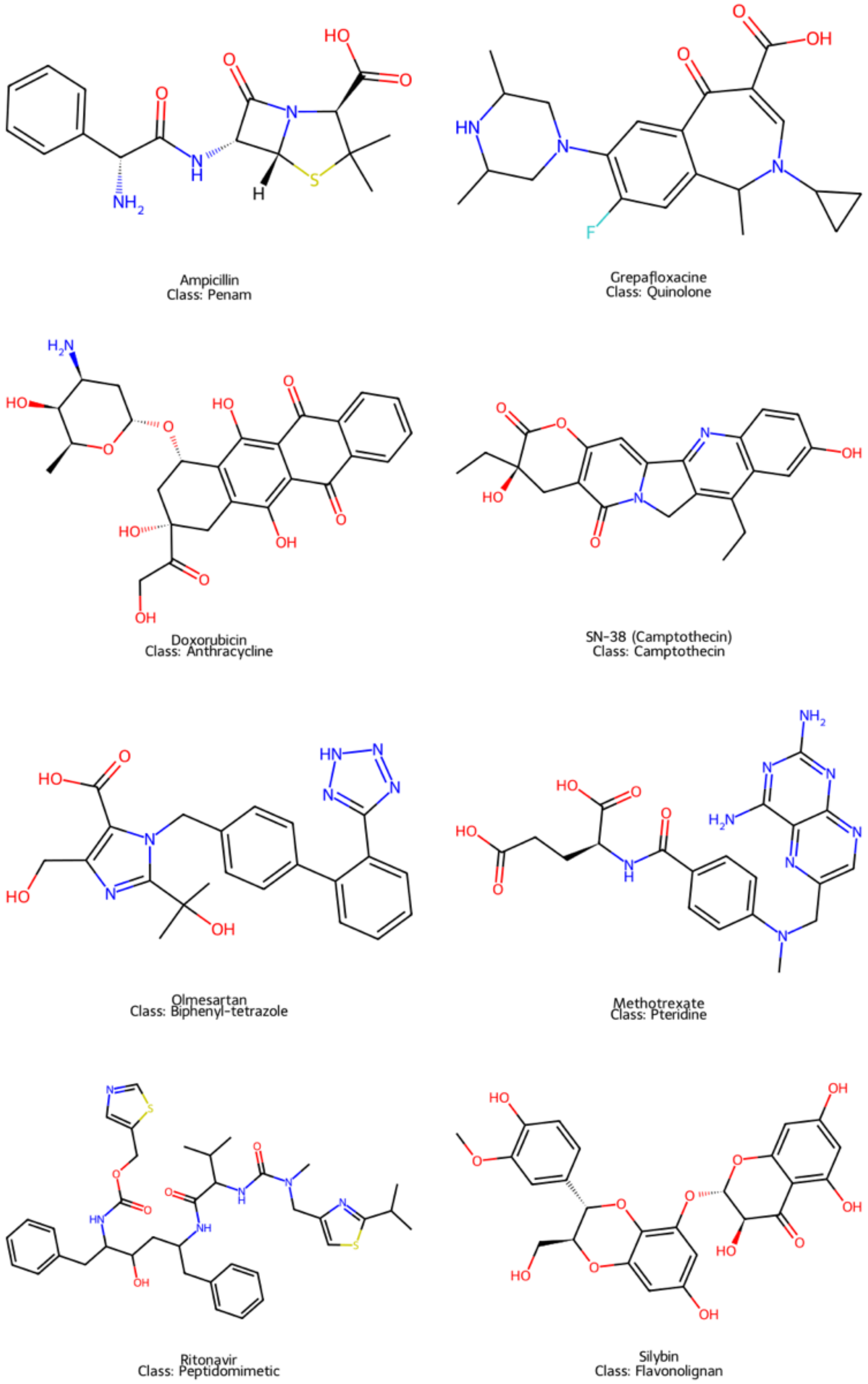


**Supplementary Figure 1.** Additional MRP2 substrate drugs. MRP2 recognises and transports structurally and physicochemically diverse classes of clinically used compounds, highlighting its broad substrate polyspecificity.

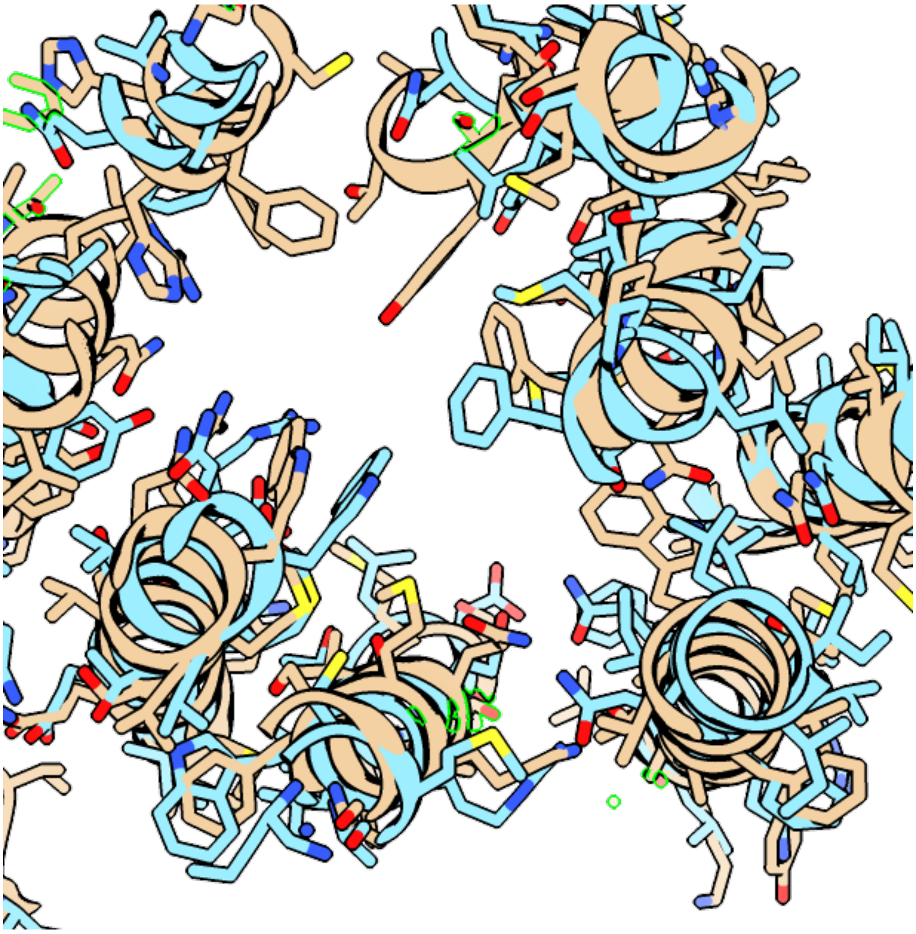

**Supplementary Figure 2.** The rMrp2 (orange) and MRP2 (cyan) share a well conserved TMD.

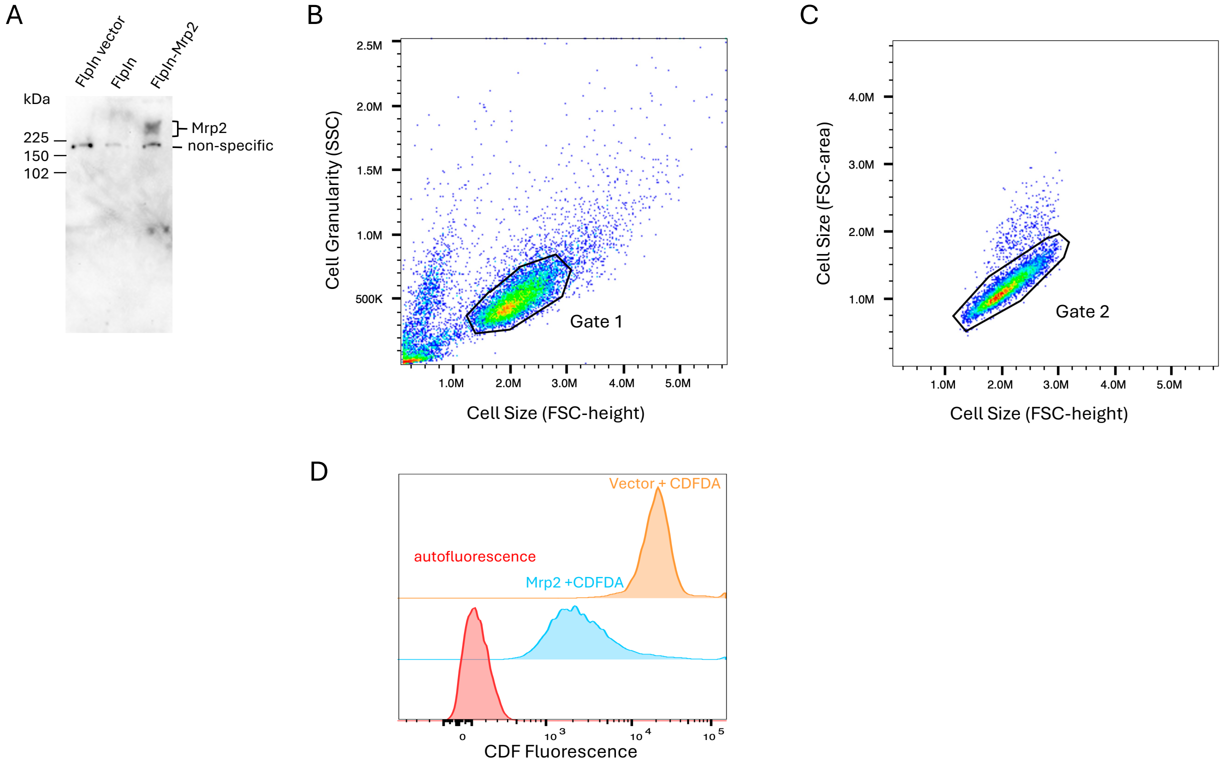


**Supplementary Figure 3**. Mrp2 is expressed and functional in the HEK293-FlpIn-rMrp2 cells. (a) Western analyses shows rMrp2 is expressed in the HEK293-FlpIn-rMrp2 cells. The primary sequence of rMrp2 encodes a protein of 173kDa which is also likely to be glycosylated resulting in a diffuse high molecular weight signal on the blot. (b) FlowJo dotplot to isolate cells of normal size and granularity (gate 1). (c) FlowJo dotplot to select single cells (gate 2). (d) Histograms showing CDF accumulation in the gated HEK293-FlpIn-rMrp2 and vector-only cells following incubation with CDFDA in comparison to the autofluorescence of untreated cells.

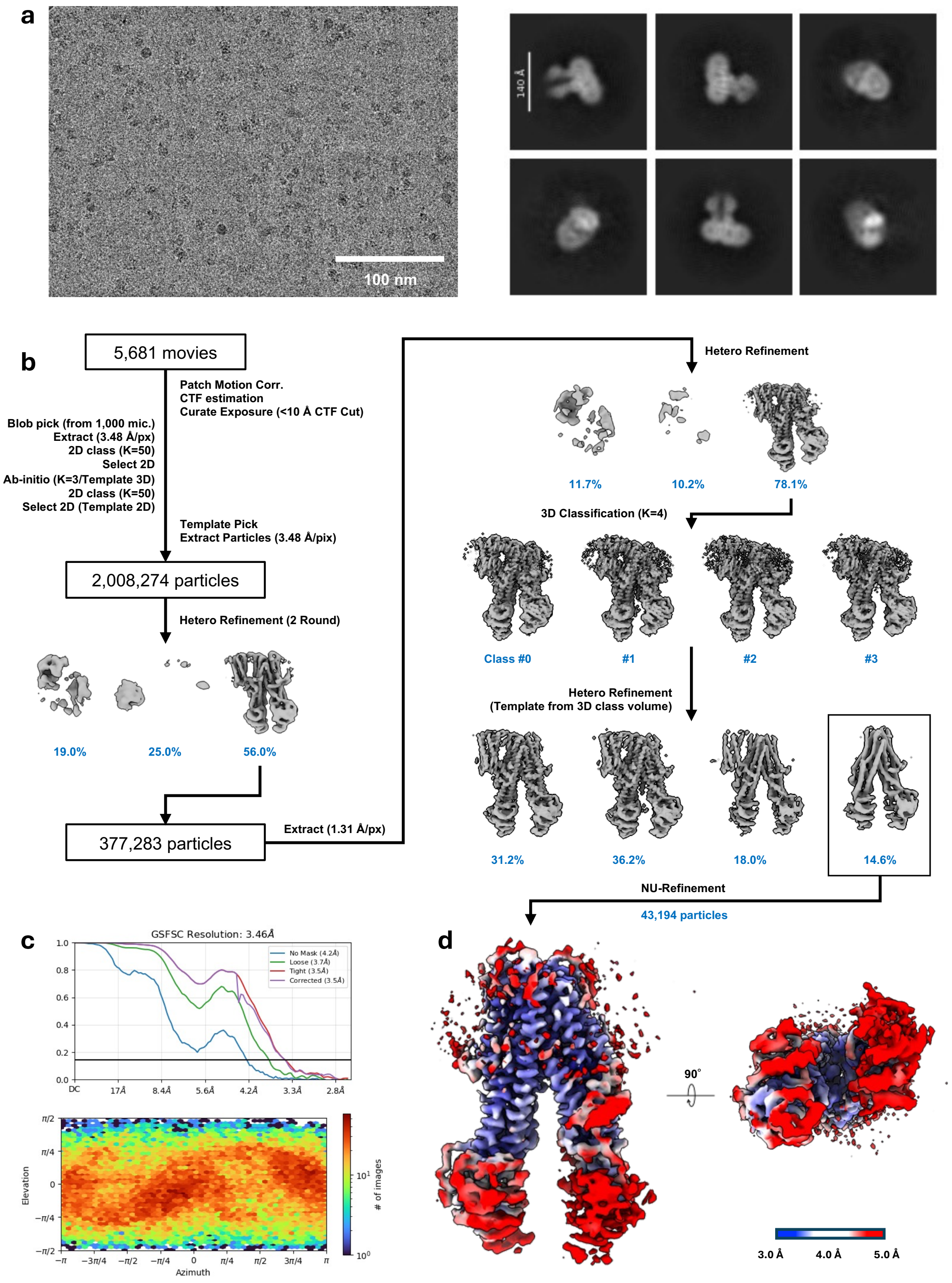


**Supplementary Figure 4.** Cryo-EM data collection and analysis workflow rMrp2 bound to vincristine. (a) Representative micrograph and selected 2D class averages. (b) Data processing workflow. (c) Gold-standard FSC curves of the final maps and Euler angle distribution plots. The resolution cut-off was FSC=0.143. (d) The maps are coloured by local resolution.

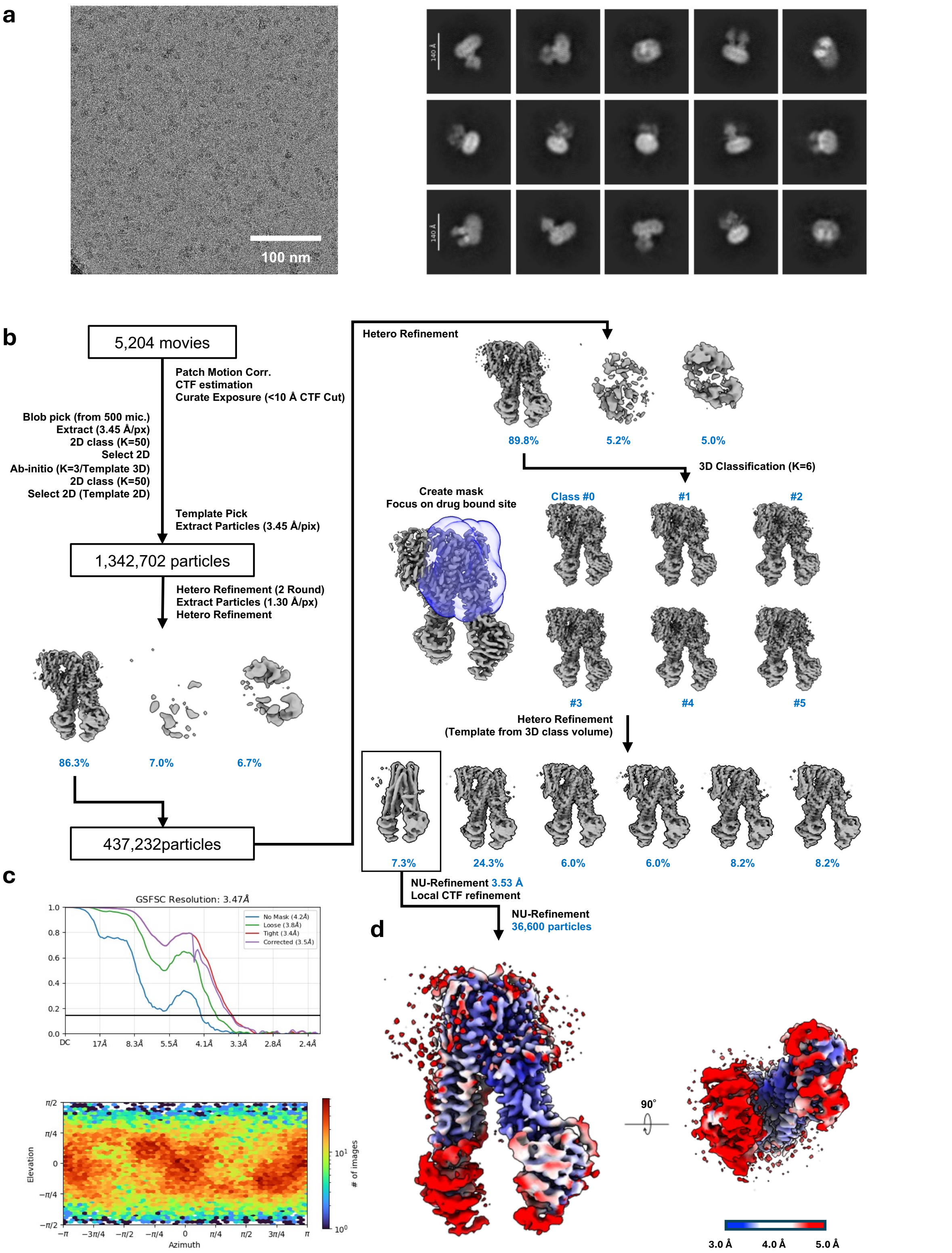


**Supplementary Figure 5.** Cryo-EM data collection and analysis workflow rMrp2 bound to vinblastine. (a) Representative micrograph and selected 2D class averages. (b) Data processing workflow. (c) Gold-standard FSC curves of the final maps and Euler angle distribution plots. The resolution cut-off was FSC=0.143. (d) The maps are coloured by local resolution.

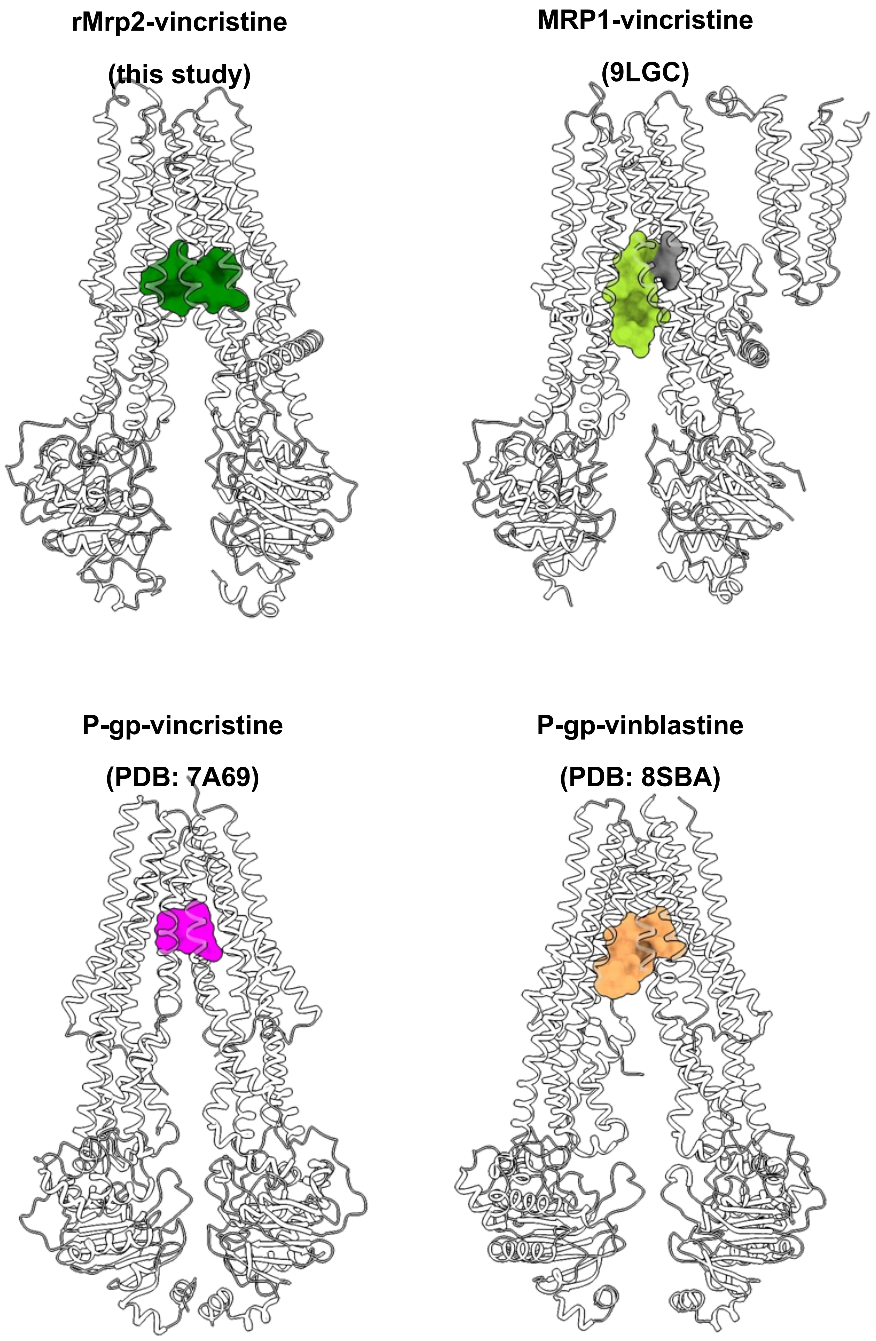


**Supplementary Figure 6.** Comparison of vinca alkaloid binding poses and stoichiometries across different ABC transporters. rMrp2 binds two vincristine molecules like MRP1 but in different poses (top panel). P-gp binds a single vincristine molecule but two vinblastine molecules (bottom panel). Neither the drug pose nor the binding binding sites are conserved between the three ABC transporters. The two vinca alkaloids are shown in surface representation and the protein in cartoon (grey); all the transporters were aligned onto the transmembrane domain and displayed in the same orientation. The associated PDB IDs are shown within the different panels.

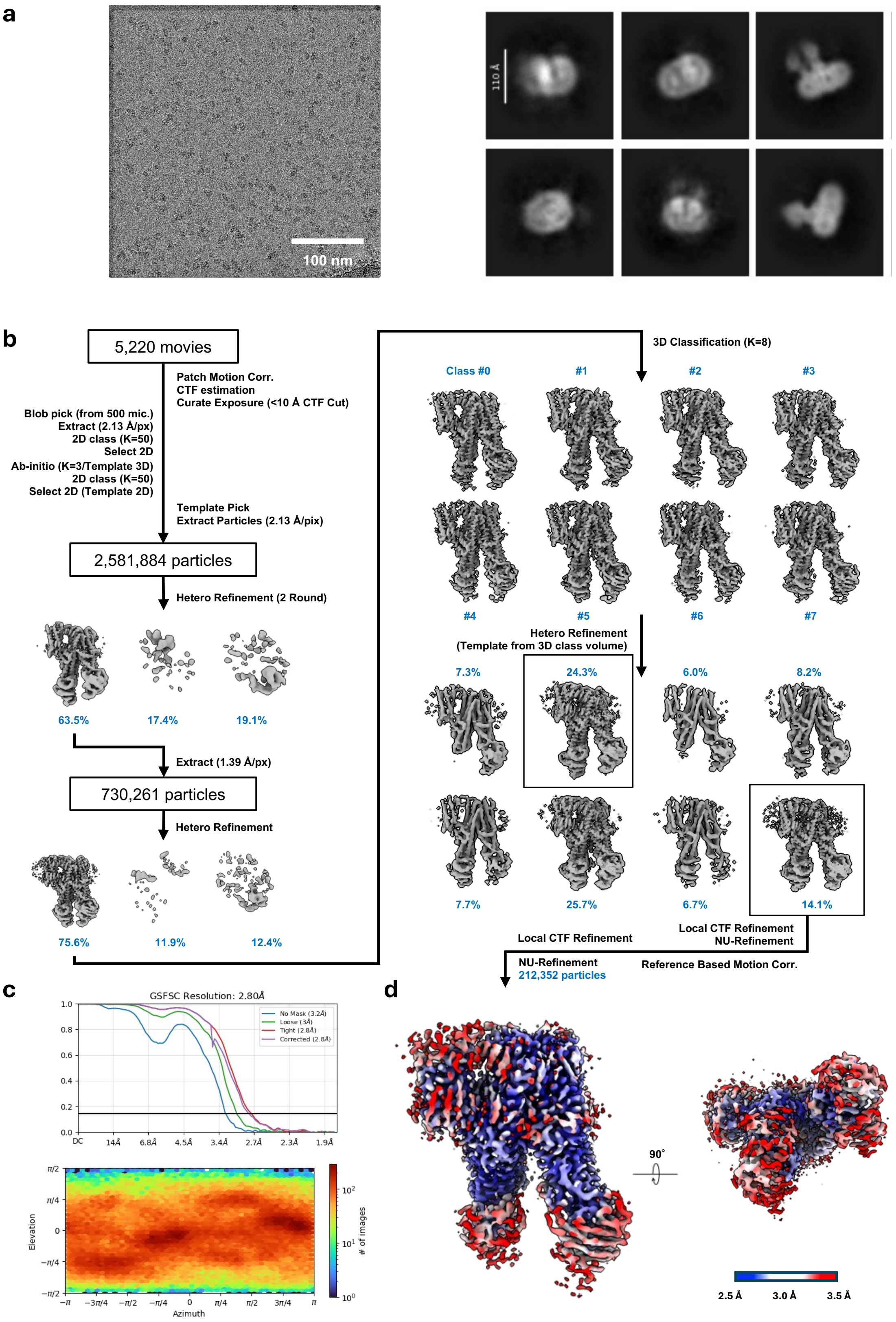


**Supplementary Figure 7.** Cryo-EM data collection and analysis workflow rMrp2 bound to teniposide. (a) Representative micrograph and selected 2D class averages. (b) Data processing workflow. (c) Gold-standard FSC curves of the final maps and Euler angle distribution plots. The resolution cut-off was FSC=0.143. (d) The maps are coloured by local resolution.

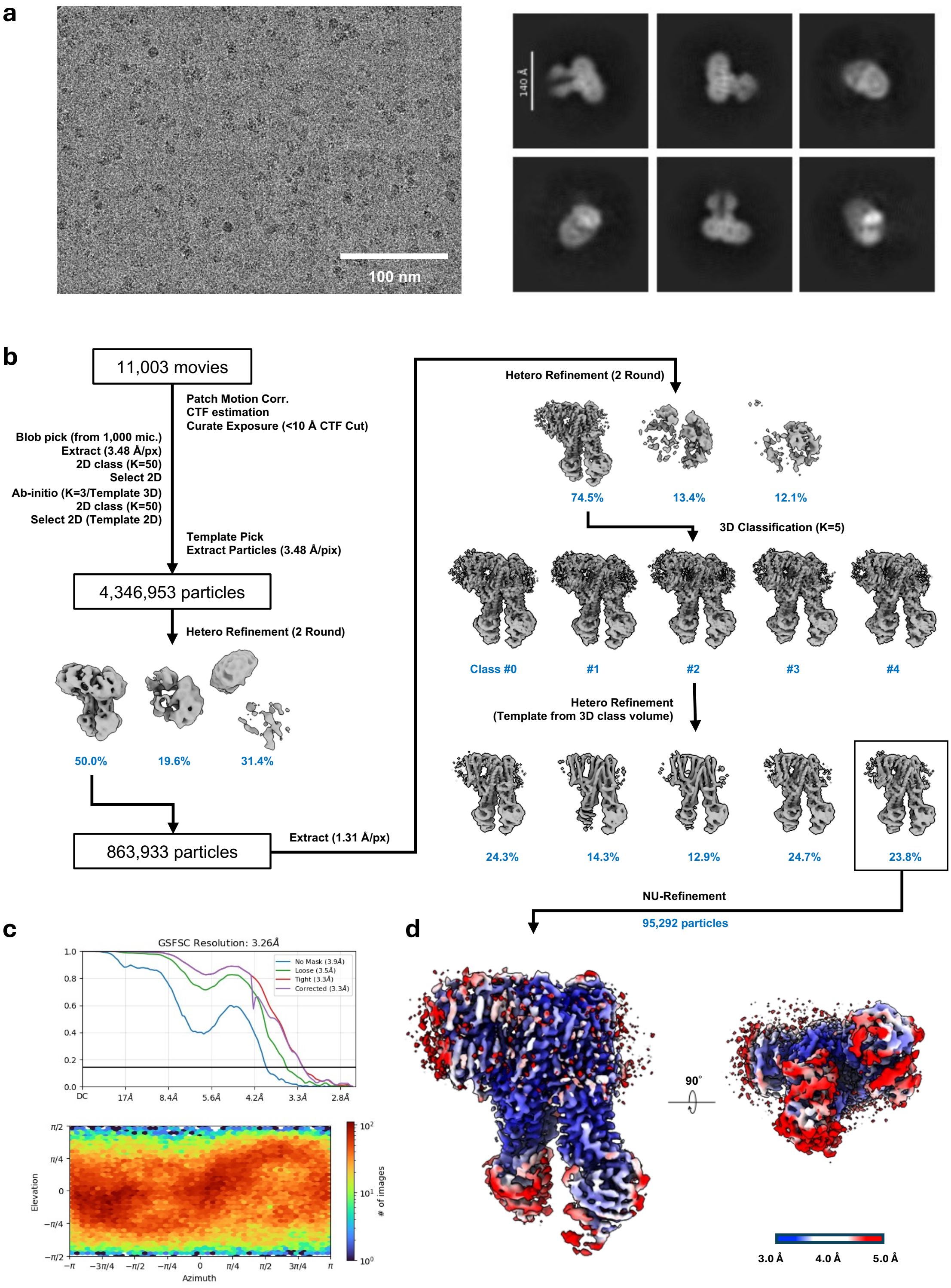


**Supplementary Figure 8.** Cryo-EM data collection and analysis workflow rMrp2 bound to sulindac. (a) Representative micrograph and selected 2D class averages. (b) Data processing workflow. (c) Gold-standard FSC curves of the final maps and Euler angle distribution plots. The resolution cut-off was FSC=0.143. (d) The maps are coloured by local resolution.

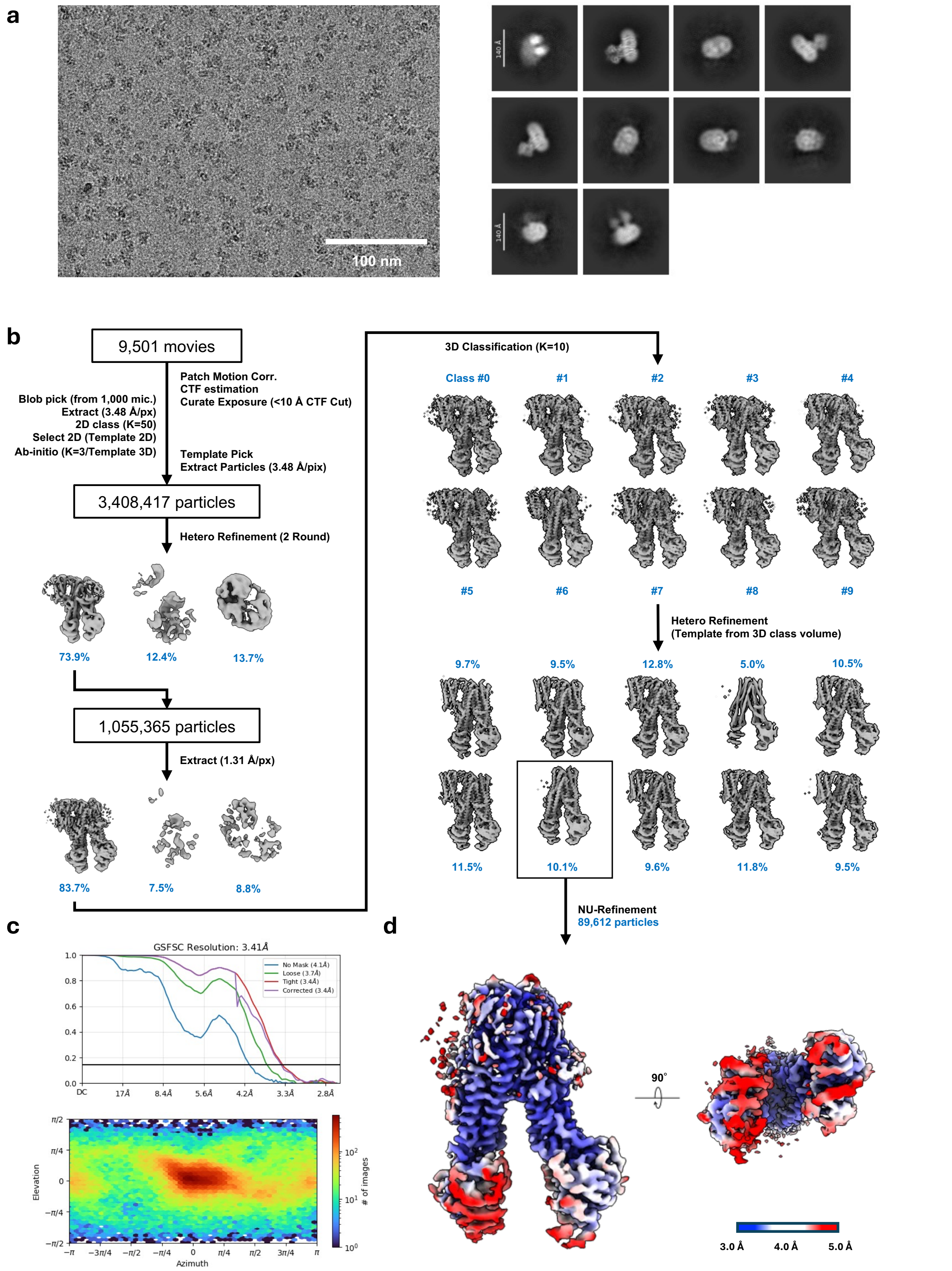


**Supplementary Figure 9.** Cryo-EM data collection and analysis workflow rMrp2 bound to bosentan. (a) Representative micrograph and selected 2D class averages. (b) Data processing workflow. (c) Gold-standard FSC curves of the final maps and Euler angle distribution plots. The resolution cut-off was FSC=0.143. (d) The maps are coloured by local resolution.

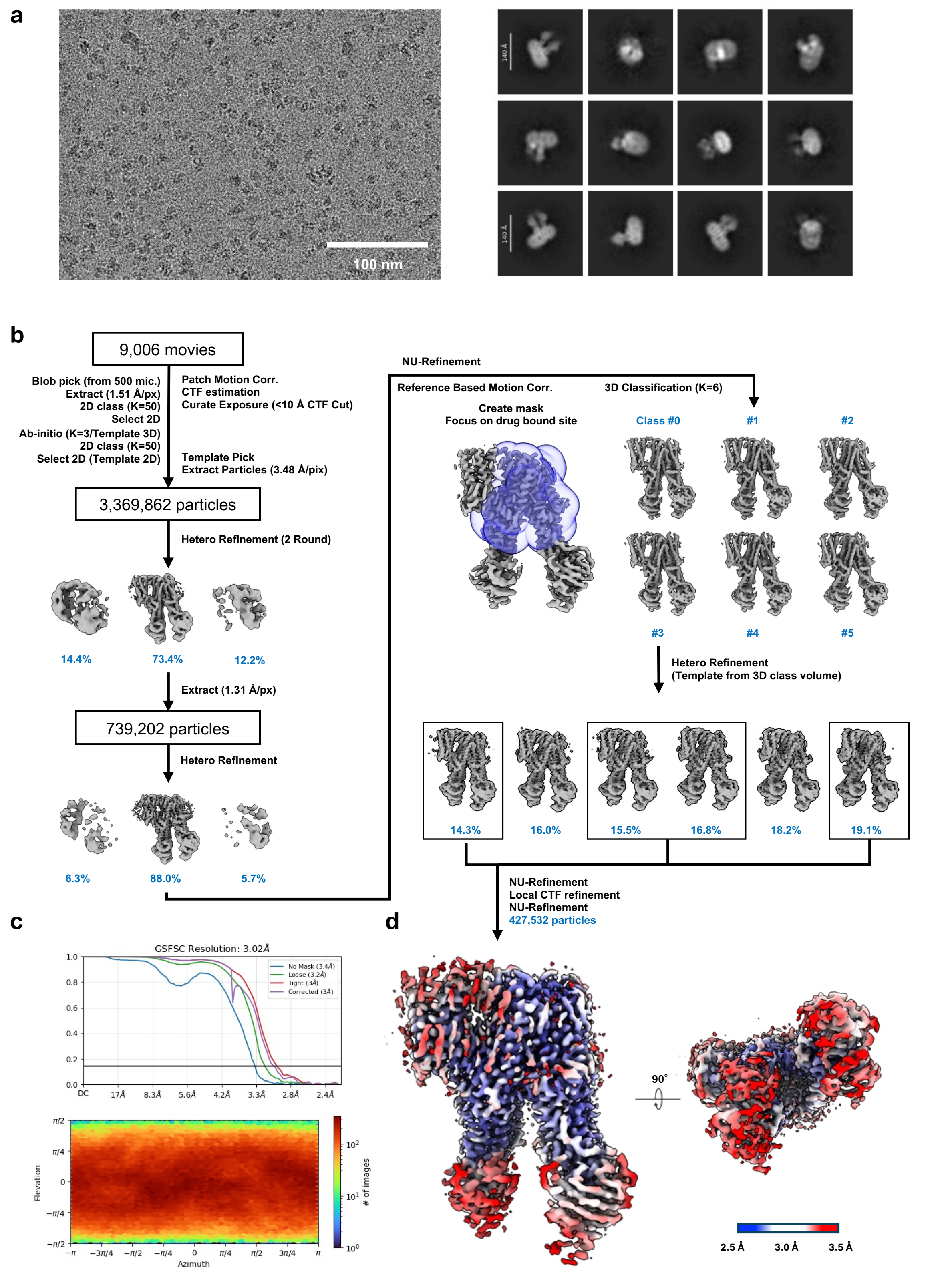


**Supplementary Figure 10.** Cryo-EM data collection and analysis workflow rMrp2 bound to MK-571. (a) Representative micrograph and selected 2D class averages. (b) Data processing workflow. (c) Gold-standard FSC curves of the final maps and Euler angle distribution plots. The resolution cut-off was FSC=0.143. (d) The maps are coloured by local resolution.

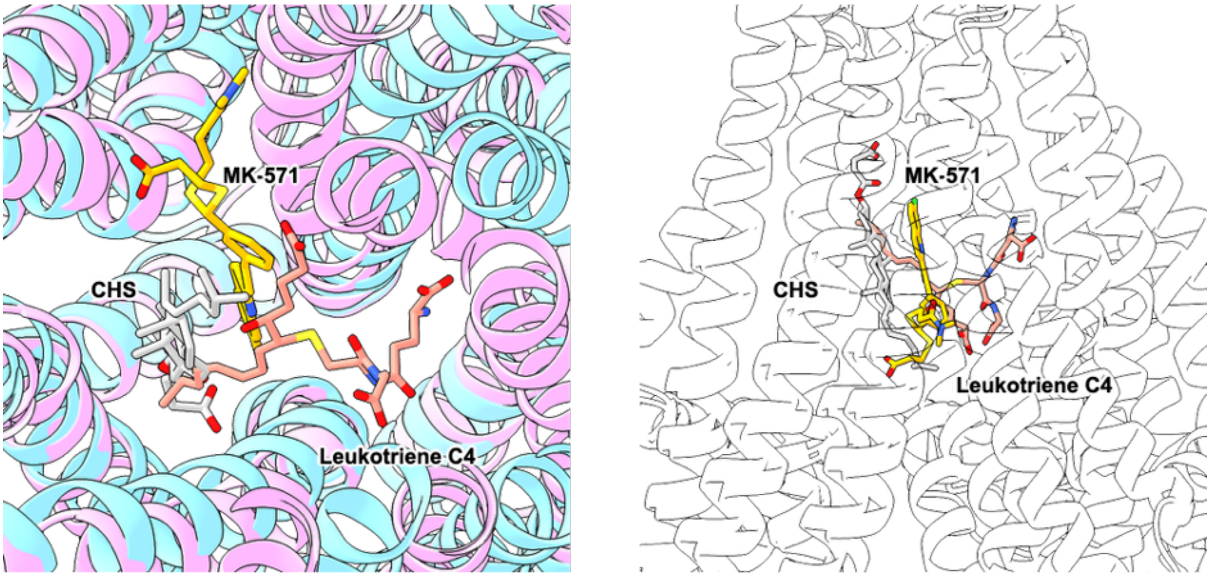


**Supplementary Figure 11.** Comparison of MK-571 and leukotriene C4 binding poses. Structures of leukotriene C4 bound to MRP2 (9C12, pink) and rMrp2 in complex with MK-571 (this study, sky blue) are shown as cartoons. Ligands are shown as sticks, with leukotriene C4 in plum, MK-571 in orange, and CHS in dark grey.

**a**

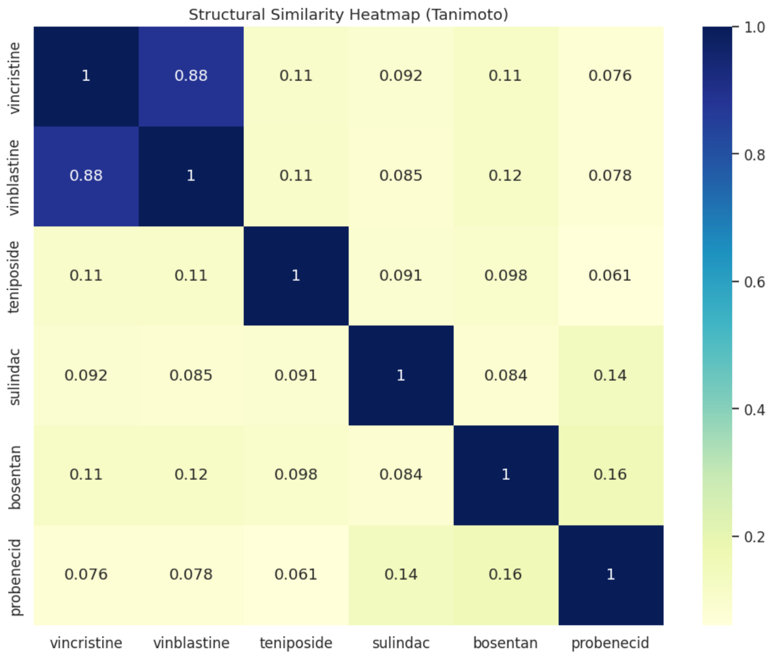


**b**

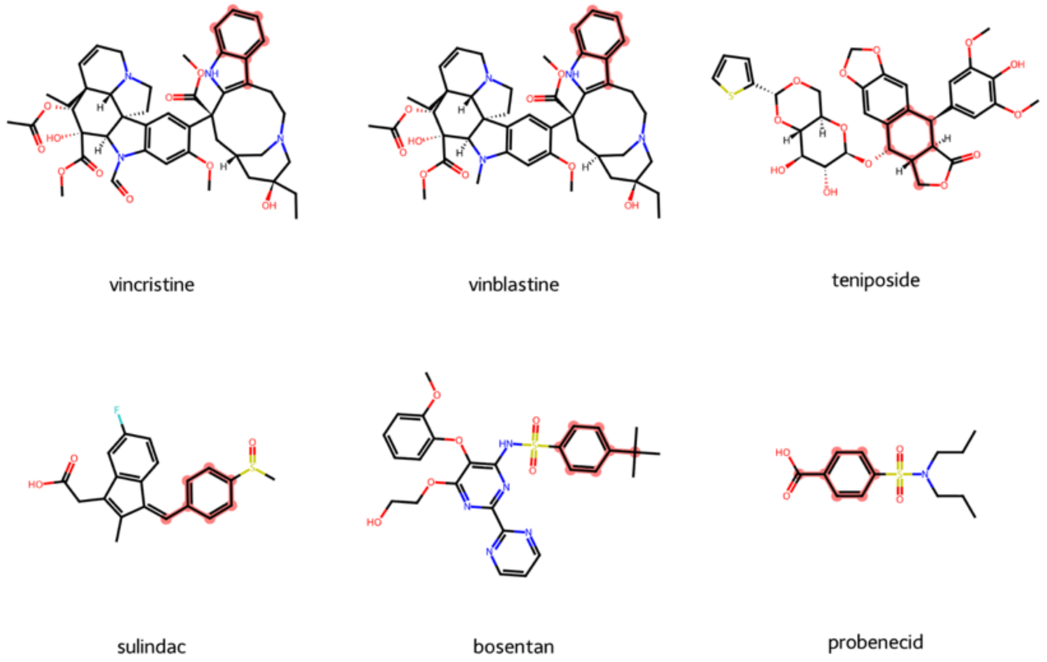


**Supplementary Figure 12.** Molecular structural similarity analysis of MRP2 drugs (this study). (a) The heatmap displays the pairwise Tanimoto similarity scores calculated using Morgan Fingerprints for the set of molecules. Values range from 0 (completely dissimilar) to 1 (identical). Higher values (darker blue) indicate greater structural similarity. The diagonal represents self-similarity (score of 1). (b) Maximum Common Substructure (MCS) analysis among the six drugs. The common substructure identified by RDKit's MCS algorithm is highlighted in red. A tolyl moiety, toluene has been identified as a common substructure in the different drug classes. Sulindac, bosentan and probenecid share a phenyl-methylsulfinyl moiety.

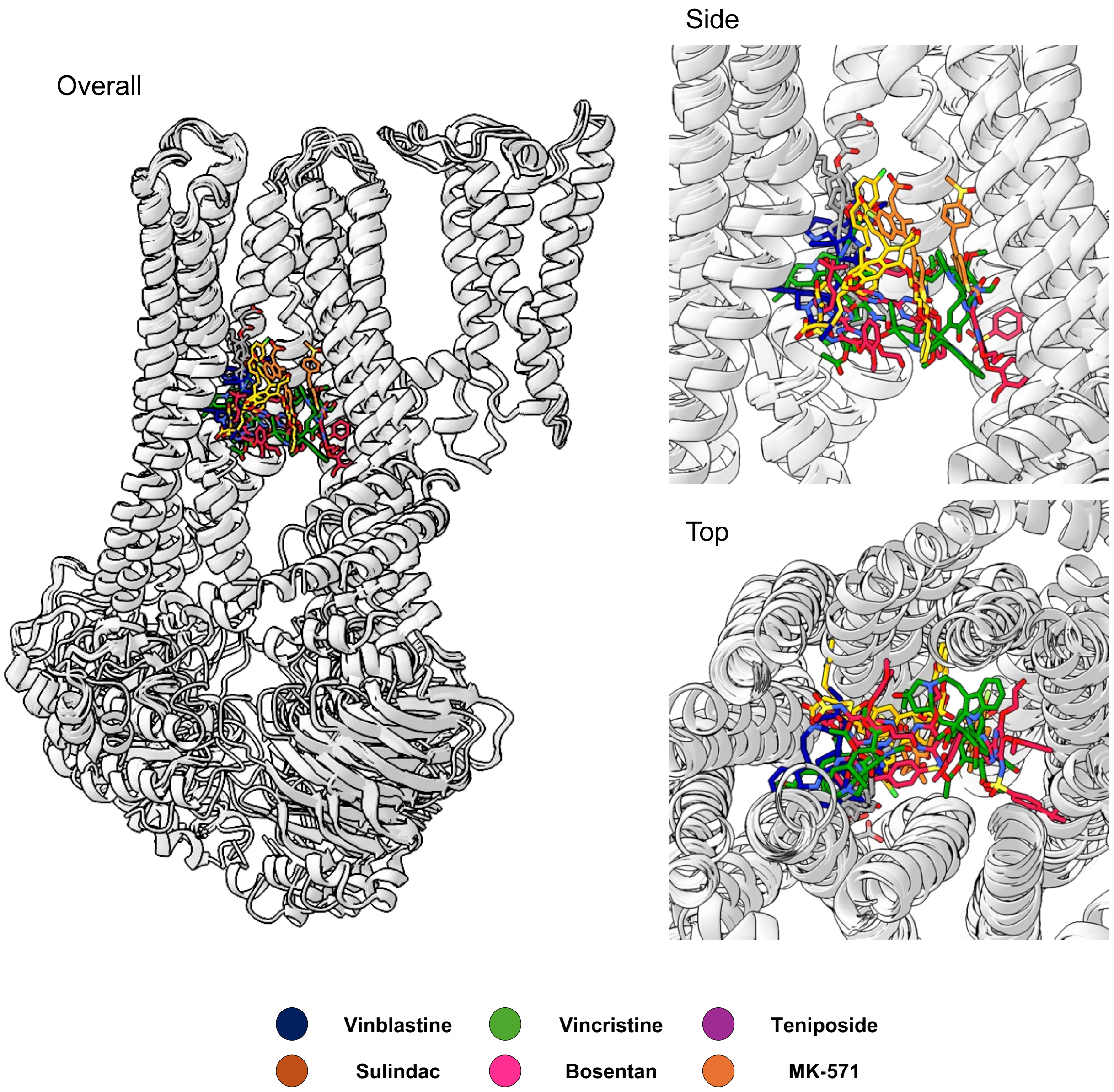


**Supplementary Figure 13.** Comparison of all the rMRP2 drug-bound structures. The structures of rMRP2 bound to each drug are superimposed. The drugs are shown in different colours, and CHS is shown in grey. In the overall view and the enlarged side view, residues 410–620 are not shown for clarity.

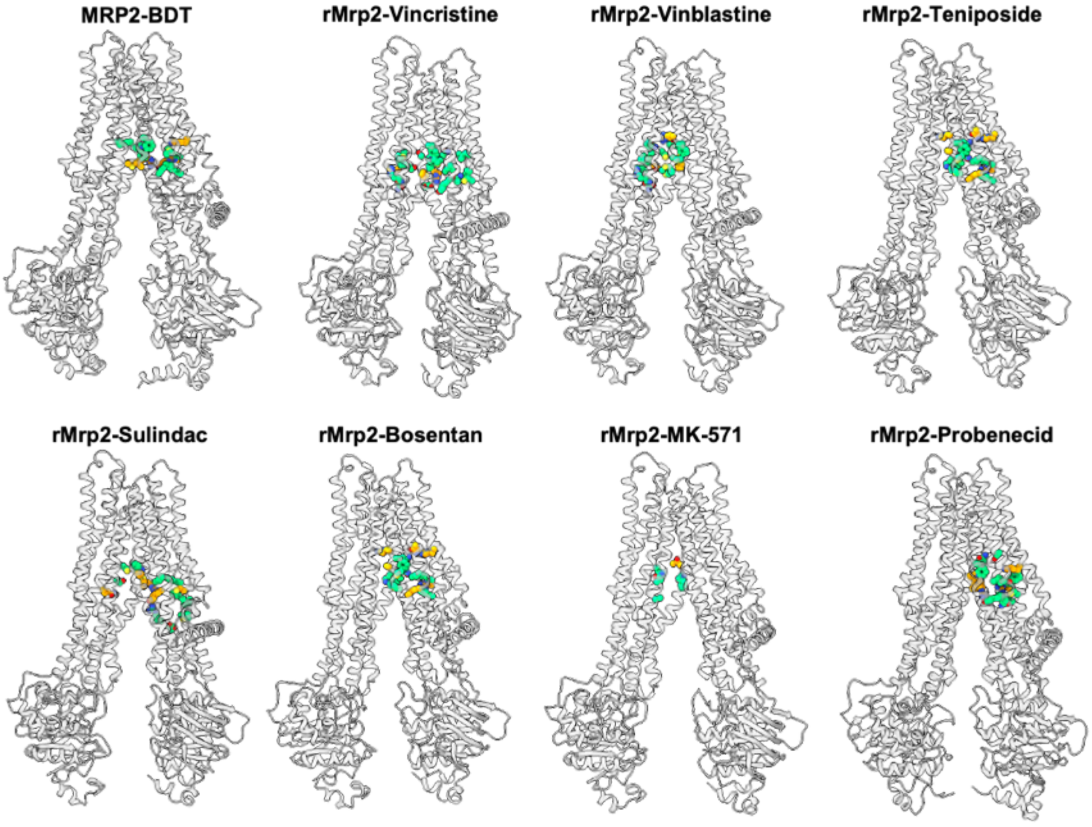


**Supplementary Figure 14.** Comparison of the ligand binding residues of rMRP2 with different drugs. The protein is shown as a grey cartoon. Side chain residues involved in ligand binding are highlighted and shown in stick model. The residues making van der Waals contacts with the ligand are coloured green and the ones forming hydrogen bonds are coloured in orange.

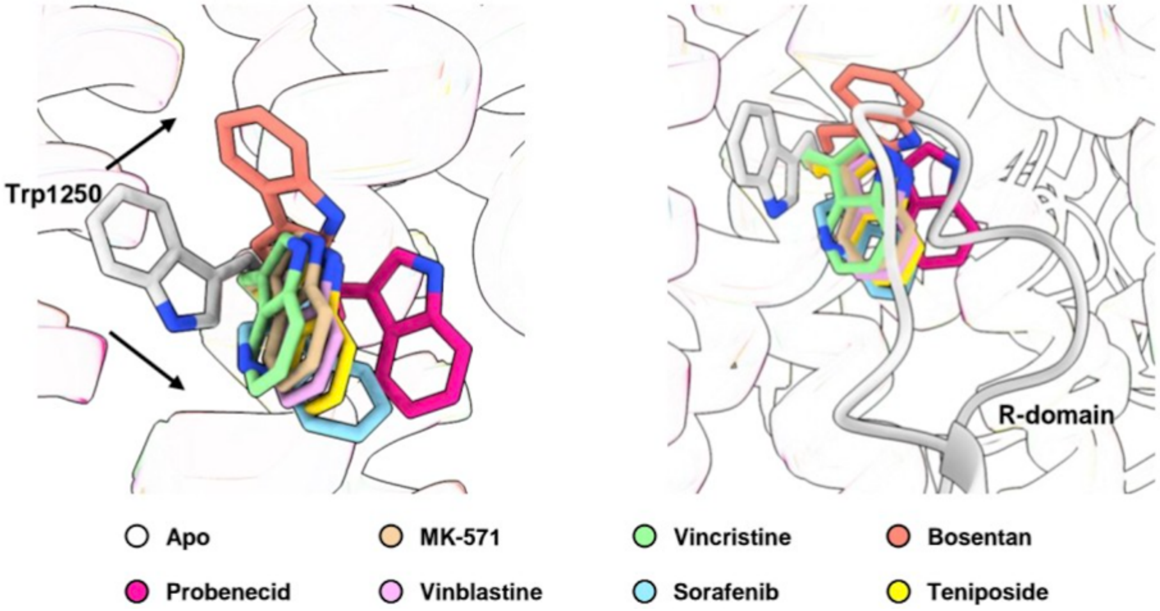


**Supplementary Figure 15.** Conformation of Trp1250 between the autoinhibited and drug-bound structures. Trp1250 is pushed away by the R-domain in the autoinhibited state, and it swings towards the TMD upon R-domain displacement.

**Supplementary Table 1.** Physicochemical properties of the clinical drugs from Figure 1a. The values were calculated using RDKit (https://www.rdkit.org/).

| Compound Name | Molecular Weight (Da) | LogP (Wildman-Crippen) | H-Bond Donors | H-Bond Acceptors | TPSA ($A^2$) |
|---|---|---|---|---|---|
| Teniposide | 656.66 | 1.15 | 3 | 14 | 165.29 |
| Vincristine | 824.93 | 2.54 | 3 | 16 | 210.12 |
| Vinblastine | 810.99 | 3.19 | 3 | 15 | 201.12 |
| Sulindac | 356.42 | 3.56 | 1 | 3 | 54.37 |
| Bosentan | 551.62 | 2.76 | 2 | 10 | 145.47 |

**Supplementary Table 2.** Pairwise comparison of $IC_{50}$ data for inhibition of CDF efflux from cells. $IC_{50}$ data are expressed as mean +/- SEM. Pairwise P-values are calculated using an unpaired, two-tailed t-test. NS = non significant.

| | MK-571 72±26 μM | Probenecid 1.5±1.2 mM | Vincristine 1.9±0.2 mM | Vinblastine 0.51±0.13 mM | Sulindac 0.62±0.17 mM |
|---|---|---|---|---|---|
| MK-571 72±26 μM | | NS | 0.0008 | 0.0258 | 0.0333 |
| Probenecid 1.5±1.2 mM | | | NS | NS | NS |
| Vincristine 1.9 ± 0.2 mM | | | | 0.0082 | 0.0082 |
| Vinblastine 0.51±0.13 mM | | | | | NS |
| Sulindac 0.62±0.17 mM | | | | | |

**Supplementary Table 3.** Primer sequences used to build pcDNA5/FRT-Mrp2 were designed in NEBuilder. Extension to the 5' end of the primers to generate overlap regions for Gibson assembly are in lower case and the engineered stop codon for rMrp2 is highlighted in red.

| Primer ID | Oligonucleotide sequence (5' to 3') |
|---|---|
| **rMrp2 forward** | tatagggagacccaagctggATGGACAAGTTCTGCAAC |
| **rMrp2 reverse** | gccctctagactcgagctcaGAGCTCTGTGTGATTCAC |
| **rMrp2 2330F** | GCCTATCAAGATGCTGACAT |
| **rMrp2 2311R** | CAGAATATAGATGTCAGCAT |
| **pcDNA5FRT forward** | tgaGCTCGAGTCTAGAGG |
| **pcDNA5FRT reverse** | CCAGCTTGGGTCTCCCTATAG |

**Supplementary Table 4.** Data collection, processing and refinement statistics for the rMrp2 in complex with the different molecules.

| | rMrp2-Vincristine | rMrp2-Vinblastine | rMrp2-Teniposide | rMrp2-Sulindac | rMrp2-Bosentan | rMrp2-MK-571 |
|---|---|---|---|---|---|---|
| **PDB entry** | 26BN | 26BO | 26BP | 26BQ | 26BR | 26BS |
| **EMDB entry** | EMD-80514 | EMD-80515 | EMD-80516 | EMD-80517 | EMD-80518 | EMD-80519 |
| **Data collection and processing** | | | | | | |
| Magnification | 130,000 | 130,000 | 130,000 | 130,000 | 130,000 | 130,000 |
| Microscope | Titan Krios | Titan Krios | Titan Krios | Titan Krios | Titan Krios | Titan Krios |
| Voltage (kV) | 300 | 300 | 300 | 300 | 300 | 300 |
| Detector | Gatan K3 | Falcon 4i | Falcon 4i | Gatan K3 | Gatan K3 | Gatan K3 |
| Energy filter | Gatan Quantum-LS, 15 eV slit | Serectris X | Serectris X | Gatan Quantum-LS, 15 eV slit | Gatan Quantum-LS, 15 eV slit | Gatan Quantum-LS, 15 eV slit |
| Electric exposure ($e^-$/Å) | 50 | 50 | 50 | 50 | 50 | 50 |
| Deforcus range (µm) | -0.8 to -1.8 | -0.8 to -1.8 | -0.8 to -1.8 | -0.8 to -1.8 | -0.8 to -1.8 | -0.8 to -1.8 |
| Pixel size (Å/px) | 0.653 | 0.921 | 0.921 | 0.653 | 0.653 | 0.653 |
| Data Processing Program | cryoSPARC (v.4.6.2) | cryoSPARC (v.4.6.2) | cryoSPARC (v.4.6.3) | cryoSPARC (v.4.6.2) | cryoSPARC (v.4.6.2) | cryoSPARC (v.4.6.2) |
| Movies | 5,681 | 5,204 | 5,220 | 11,003 | 9,501 | 9,006 |
| Initial / Final particle images (no | 2,008,274 / 43,194 | 1,342,702 / 36,600 | 2,581,884 / 212,352 | 4,346,953 / 95,292 | 3,408,417 / 89,612 | 3,369,862 / 427,532 |
| Symmetry imposed | C1 | C1 | C1 | C1 | C1 | C1 |
| Map resolution (Å) | 3.46 | 3.47 | 2.80 | 3.26 | 3.41 | 3.02 |
| FSC threshold | 0.143 | 0.143 | 0.143 | 0.143 | 0.143 | 0.143 |
| **Refinement** | | | | | | |
| Refinement Program | PHENIX (v.1.20.1) | PHENIX (v.1.20.1) | PHENIX (v.1.20.2) | PHENIX (v.1.20.1) | PHENIX (v.1.20.1) | PHENIX (v.1.20.1) |
| Model resolution (Å) | 3.39 | 3.41 | 2.76 | 3.20 | 3.32 | 2.96 |
| FSC threshold | 0.143 | 0.143 | 0.143 | 0.143 | 0.143 | 0.143 |
| Model composition | | | | | | |
| Non-hydrogen atoms | 9,247 | 9,179 | 11,107 | 11,083 | 9,244 | 11,094 |
| Protein residues | 1,157 | 1,156 | 1,389 | 1,390 | 1,157 | 1,389 |
| Ligands | 2 Vincrastine (VNC) | 1 Vinblastine (VLB) | 1 Teniposide (9TP)<br>1 CHS (Y01) | 2 Sulindac (SUZ) | 3 Bosentan (K86) | 1 MK-571 (R)<br>1 CHS (Y01) |
| R.m.s. deviations | | | | | | |
| Bond length (Å) | 0.003 | 0.002 | 0.003 | 0.003 | 0.003 | 0.003 |
| Bond angles (°) | 0.843 | 0.734 | 0.548 | 0.638 | 0.576 | 0.568 |
| Validation | | | | | | |
| MolProbity score | 1.73 | 1.89 | 2.23 | 1.78 | 1.89 | 1.74 |
| Clashscore | 7.83 | 10.47 | 8.76 | 7.75 | 8.67 | 7.35 |
| Ramachandran plot | | | | | | |
| Favored / Allowed (%) | 95.66 / 4.00 | 94.87 / 4.96 | 95.59 / 4.05 | 94.87 / 4.84 | 93.66 / 5.91 | 95.23 / 4.56 |
| Disallowed (%) | 0.35 | 0.17 | 0.36 | 0.29 | 0.43 | 0.22 |
| Mask CC | 0.82 | 0.82 | 0.79 | 0.82 | 0.83 | 0.84 |
